# Validated phytohormone analysis with paired untargeted workflow enables new insights into soil phytohormonome

**DOI:** 10.64898/2026.03.20.713310

**Authors:** Sarah L. Lane, Rashaduz Zaman, James F. Cahill, Carolyn J. Fitzsimmons, Lauren A.E. Erland

## Abstract

The contribution of soil chemistry to plant growth and resilience, including presence of phytohormones, is increasingly recognized, yet characterization is limited by chemical complexity of soil matrices, diversity and low-abundance of metabolites. To enable further discoveries, we developed and characterized performance of a liquid chromatography-mass spectrometry method with solid phase extraction, integrating targeted and untargeted hormonomic approaches for comprehensive soil phytohormone profiling. Method performance was evaluated for fifteen plant growth–regulating compounds and precursors, including abscisic, gibberellic, jasmonic, and salicylic acids, auxins, cytokinins, karrikins, melatonin, and tryptophan, showing strong linearity (R² = 0.989–0.999), sensitivity (limits of detection and quantification 0.1–50.2 and 1.4–167.3 pg on-column, respectively), and precision (1.3–9.6% intraday; 3.4–34.8% interday). Soil composition impacted recovery; however, for most phytohormones rates were within 20% of matrix-adjusted spiked value, showing robustness across sandy, peat-rich, and clay-rich textures and suitable for use. We used the method to quantify analytes in research-relevant, active soils. Integration of untargeted analysis expanded coverage to 250 additional putative phytohormones and related metabolites, revealing chemical signatures potentially associated with plant community composition. This approach provides a versatile framework for investigating belowground phytohormone dynamics and their roles in plant physiology, resilience, and soil–plant feedbacks.

## Introduction

Soil metabolomics is an emerging discipline focused on uncovering the diverse network of soil chemistry underpinning plant-soil interactions, which can help us to start to answer questions like “What is deposited there?”, “By what?” and “Why?”. Plant-soil interactions are emerging as key factors governing plant resilience to changing landscapes at the population and community level, and these interactions occur through a wide range of metabolites including lipids, sugars, protein-incorporating and non-protein amino acids, volatile organic chemicals, terpenoids and phenylpropanoids (Das et al., 2016; Ma et al., 2022; McLaughlin et al., 2023; Upadhyay et al., 2022; Vives-Peris et al., 2017). Phytohormones are one group of metabolites present in soils, and are likely contributors to plant-plant and plant-microbiome dynamics including allelopathy (limiting growth of neighboring plants), recruitment of nitrogen-fixing bacteria (such as in alder or legumes), and recruitment of mycorrhizal fungal partners (Gueddou et al., 2022; Hickman et al., 2025; Li et al., 2025). Phytohormones classically include auxins, cytokinins, gibberellin, abscisic acid (ABA) and ethylene, and have been expanded to include several groups of compounds, many of which were originally recognized as specialized metabolites including jasmonic acid (JA), brassinosteroids, salicylic acid (SA), strigolactones, indoleamines and karrikins (Omoarelojie et al., 2019; Wang and Irving, 2011). Sources of hormones in soils extend beyond the presence of plant materials or root exudates, and extend to plant growth-promoting bacteria and other microorganisms that produce or regulate phytohormones and their derivatives to induce growth effects or protective effects and further contribute to the complexity of the soil phytohormonome. While above ground plant-plant communication through production of volatile hormones (methyl salicylate and methyl jasmonate) and secondary metabolites is well established, considerably less is understood underground (Karban et al., 2014; Sakurai and Ishizaki, 2024). More research is needed to elucidate soil phytohormonome composition and dynamics, including how phytohormones mediate plant-soil interactions.

Hormonomics is the comprehensive study of phytohormone precursors, metabolites, conjugates and derivatives, enabling both qualitative and quantitative analysis of this complex and dynamic system (Šimura et al., 2018). However, a major challenge to converting this tool for use in soil metabolomics is the highly complex matrix of soil components that can cause significant interference when extracting chemical compounds (Song et al., 2024). Typical soil extractions use solvents that are not appropriate for mass-spectrometry (e.g., causing ion suppression), and can alter metabolites present (e.g., causing polymerization), limiting capture of transient, fragile, or reactive metabolites. Phytohormones present an even greater quantitative challenge. While often present at nM – μM concentrations in plant tissues, phytohormones are even more dilute when present as exudates in soils, in pM – nM concentrations (Boutet-Mercey et al., 2018; Tawaraya et al., 2014, 2013). Phytohormones also represent significant chemical diversity spanning a wide range of pKa, solubility and structure. Adding complexity, phytohormones are inherently transient; their production and activity are tightly controlled through regulation of their precursors, activation or inactivation via conjugation to sugars or other moieties, and spatial or temporal separation. This limits the use of classic extraction methods that prioritize efficient recovery of very few analytes over broad analyte capture, and also those that successfully capture a diverse range of analytes (e.g. pesticides) but require conditions like heating or vacuum steps that cause degradation (Dąbrowska et al., 2003; Ibáñez et al., 1998; Lee et al., 2025).

Here we describe a validated hormonomic method which includes initial solid phase extraction followed by separation and detection using ultra-performance liquid chromatography quadrupole time-of-flight tandem mass-spectrometry (UPLC-qTOF-MS/MS) method for quantification of phytohormones from soils (Figure 1). This targeted quantification method includes ABA, IAA, indole-3-butyric acid (IBA), JA, Karrikin 1, 2, 3, 4 (KAR1 – 4), melatonin (Mel), N6-benzyladenine (BA), N6-isopentenyladenine (2IP), tryptophan (Trp), zeatin (Zea), gibberellic acid 3 (GA3) and SA, representing key representative molecules from each of the major hormone classes and several environmental plant growth regulators (PGR). This is paired with a semi-quantitative untargeted method which expands the method to include 250 hormones, their precursors, metabolites and conjugates, while also capturing a snapshot of the overall chemical space within soil. While several other methods have been described for extraction and quantification of hormones from soils, they require potentially degrading pretreatment steps such as air or vacuum drying, specialized, complicated or lengthy extraction techniques, and heated extraction steps that are not conducive to capturing less stable or volatile phytohormones (Dong et al., 2012; Lu et al., 2022). Our method overcomes these limitations by flexibly extracting and analyzing a broad spectrum of plant phytohormones from fresh-frozen soil using a combined targeted-untargeted approach, allowing a more comprehensive semi-quantitative exploration of soil-based phytohormone profiles.

**Figure 1.**
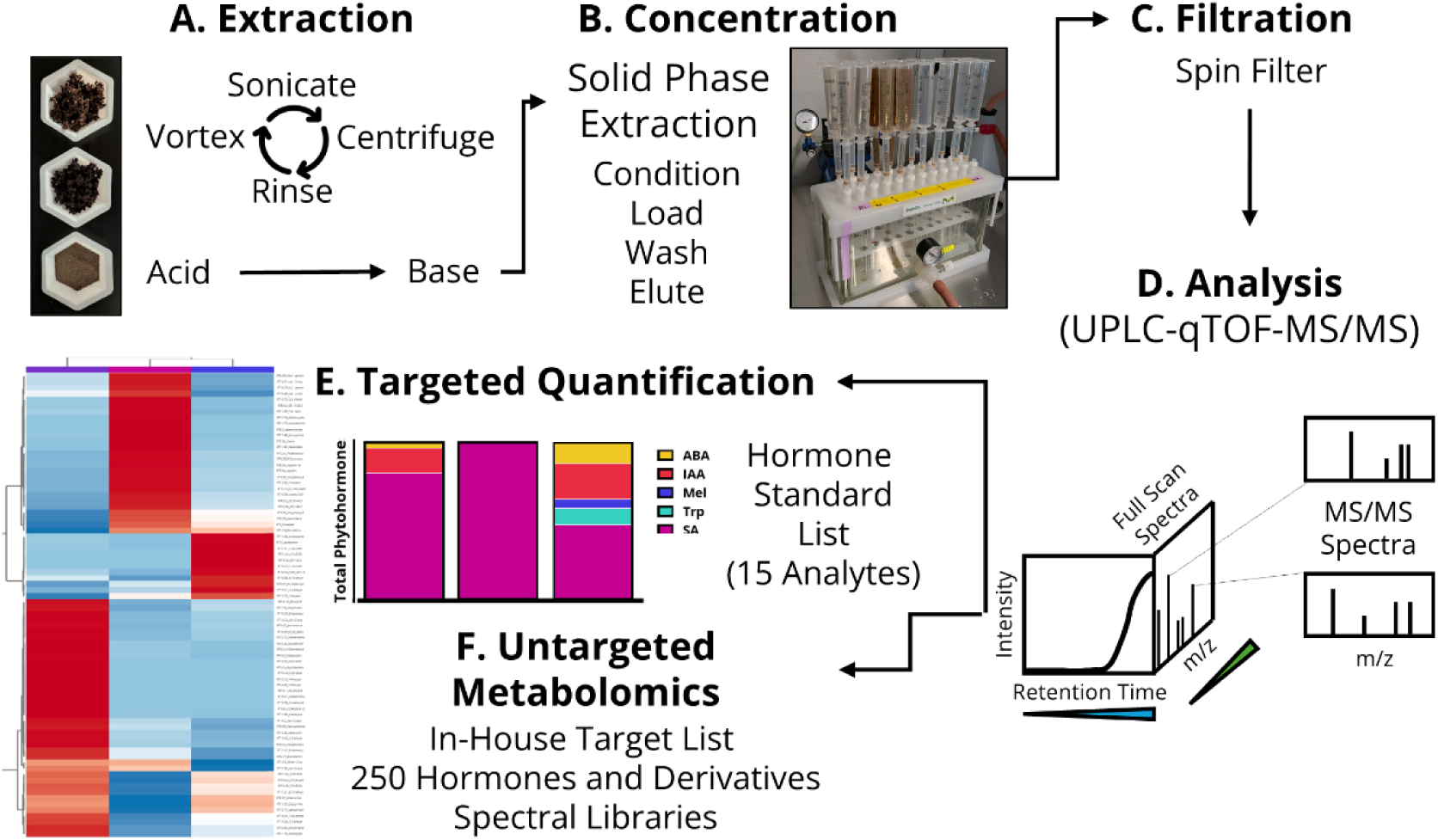
Overview of soil extraction and analysis workflow. **A.** Extraction is performed using 50% MeOH in 0.2% formic acid, followed by a repeat extraction in 0.01M NaOH. Extracts are then pooled and concentrated (**B**) using solid phase extraction with Waters Oasis® HLB cartridges to isolate phytohormones and small molecular weight compounds. Eluent is filtered to remove any remaining particulate (**C**) and analysed using ultra-high performance liquid chromatography coupled with quadrupole time-of-flight tandem mass spectrometry (**D**). Collected spectra (full scan and MS/MS) are then exported and processed. Data are analysed for quantification of 15 analytes representing a range of phytohormones and plant growth regulators across several classes (**E**). Finally, data are analysed using an untargeted process (**F**) and compared to an in-house target list containing 250 hormones and their derivatives, as well as publicly available spectral libraries for further annotation. Alt-text: A flow chart summarizing the main steps of the method: extraction, concentration, filtration, analysis, targeted quantification and untargeted metabolomics, with a bar chart of specific phytohormones and a heat map of annotated mass features as the final output.

## Methods

### Chemicals and Materials

Methanol (LC-MS grade, A452SK), acetonitrile (LC-MS grade, A995), formic acid (LC-MS grade, A117), melatonin, and indole-3-butyric acid were purchased from Fisher Scientific. Acetic acid (Suprapure) was purchased from Supelco. Sodium hydroxide, jasmonic acid, tryptophan, gibberellic acid, and salicylic acid were purchased from Sigma Aldrich. (+) abscisic acid was purchased from Millipore Sigma. 6-benzylaminopurine was purchased from Phytotech Labs. Indole-3-acetic acid, Karrikinolide (Karrikin 1), Karrikin 2, N6-(Δ2-isopentenyl)adenine, and zeatin were purchased from Caymen Chemicals. 3,5-dimethyl 2H-Furo[2,3-c]pyran-2-one (Karrikin 3) and 3,7-dimethyl 2H-Furo[2,3-c]pyran-2-one (Karrikin 4) were purchased from Toronto Research Chemicals. Ultra-pure water (18 MΩ) was acquired using a Barnstead Smart2Pure Pro 16 UV/UF Water Purification System (Thermofisher).

### Design of the Method Optimization and Validation

Our method was optimized prior to validation. We initially evaluated the effect of soil volume (1 – 10 mL), pH (acid, base, or sequential extraction), solvent volume (1 – 25 mL), SPE sorbent mass (60 or 400 mg), and elution volume (0.5 – 2.5 mL) on extractability and recovery for a subset of phytohormones and plant growth regulators. A Youden trial for main factor effects utilizing an 8-run, 6 factor fractional factorial design was performed (Karageorgou and Samanidou, 2014) to further optimize the extraction and recovery of analytes of interest. The Youden trial was conducted in agricultural field samples with a clay-rich texture spiked with 50 ng of a standard mix containing ABA, IAA, IBA, JA, KAR1, KAR2, KAR3, KAR4, Mel, BA, 2IP, Trp, Zea, GA, and SA. This concentration was selected so that full spike recovery signal would be within the linear range for all analytes after endogenous analyte contributions. Factors evaluated were prewetting (5 mL NaCl), salt content (inclusion of 1M NaCl during base extraction), efficiency of solvent contact (weight of extraction time on vortex and sonication steps), acid choice (acetic or formic acid), and order of extraction (acid or base first, Table 1). Calculated mean abundance per gram moisture-corrected dry weight for each factor are presented in Table S1 and plotted as coordinates in GraphPad Prism 10.

**Table 1.** Experimental design for Youden trial (8-run, 6 factor), fractional factorial design for 6 main factors (A: acid selection, B: salt addition, C: base strength, D: extraction order, E: agitation/contact time, F: prewetting). Three replicates were analysed for each run from pooled agricultural field soils.

| **Run** | **A/a** | **B/b** | **C/c** | **D/d** | **E/e** | **F/f** |
| --- | --- | --- | --- | --- | --- | --- |
| 1 | Acetic acid | - | 0.1N NaOH | acid, base | vortex weighted | prewet, 1 M NaCl |
| 2 | Formic acid | - | 0.1N NaOH | base, acid | sonication weighted | prewet, 1 M NaCl |
| 3 | Acetic acid | 1 M NaCl | 0.1N NaOH | base, acid | vortex weighted | - |
| 4 | Formic acid | 1 M NaCl | 0.1N NaOH | acid, base | sonication weighted | - |
| 5 | Acetic acid | - | 0.01N NaOH | acid, base | sonication weighted | - |
| 6 | Formic acid | - | 0.01N NaOH | base, acid | vortex weighted | - |
| 7 | Acetic acid | 1 M NaCl | 0.01N NaOH | base, acid | sonication weighted | prewet, 1 M NaCl |
| 8 | Formic acid | 1 M NaCl | 0.01N NaOH | acid, base | vortex weighted | prewet, 1 M NaCl |

To validate our method, we evaluated accuracy, precision, selectivity, linearity, limit of detection and quantitation. Selectivity was determined by optimizing solid phase extraction with blanks spiked with analytes of interest to ensure their capture and effective elution. Selectivity was achieved through quantitative time-of-flight tandem mass spectrometry that is compared to an in-house standard library, scheduled precursor list, and standards to ensure accurate identification of target analytes and resolution of analytes. The method was evaluated for accuracy and precision for quantification of all 15 analytes. To determine accuracy, soil samples were spiked with 50 ng of the same standard mix, then extracted using the method. Analytes were quantified, then recovery rate was established using equation 1, where C_Endo_ represents endogenous concentration of the analyte in a particular soil, and C_Measured_ is the concentration quantified in the sample. Endogenous concentrations of phytohormones in each soil type were considered in the equation to account for the effect of baseline phytohormone levels when calculating recovery rates.

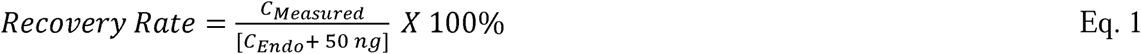

Ion suppression due to matrix effects was established through the post-extraction spike method by spiking samples for each soil type with 50 ng of the same standard mix used to establish recovery rate after extraction and prior to analysis (McManus et al., 2019). Matrix-corrected recovery rates were calculated using equations 2 and 3.

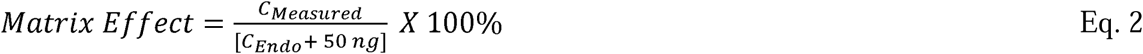

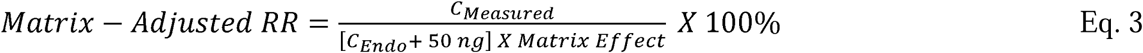

Intra- and inter-day precision was determined by calculating relative standard deviation using three replicate injections (intra) over three days (inter) that were at least two days apart. The instrument limit of detection and quantitation were determined using the standard method established by the International Conference of Harmonization with equations 4 – 5 where σ is the standard deviation for the response in the solvent blank, and S is the calibration curve slope for the analyte (Abraham, 2010). This was performed on at least 8 blank replicates collected over three weeks. Inter- and intra-day precision was calculated using relative standard deviation (RSD %) at 50 ng/mL, where SD indicates the standard deviation of the response (eq. 6).

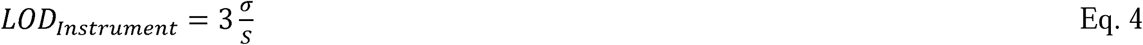

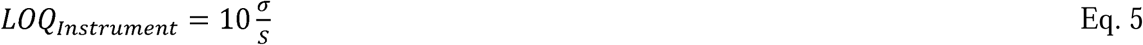

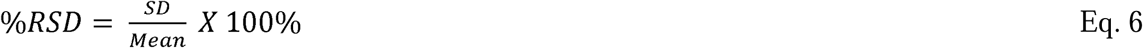

The range of quantitation was determined using a 12-point standard curve over the range from 0.0015 – 750 ug mL^-1^. Linearity of the response was calculated using R^2^ value on three replicate injections from the same day.

### Sample Collection and Preparation

Three soil types were used to optimize and characterize performance of the method. We selected non-sterile soils exposed to actively growing plants in conditions commonly encountered in plant biology, horticultural and agricultural research: a common peat-enriched greenhouse potting mix (greenhouse), clay-rich field soils with a history of agricultural cultivation (field), and sandy field soils from an undisturbed dry mixed grassland site (grassland). Soils were collected from active, ongoing studies. Greenhouse soils were collected from three-month-old Albion strawberries (*Fragaria x ananassa* ‘Albion’) grown in Sunshine Mix No. 4, then all soil from each pot was collected and preserved at -20 °C for analysis. Agricultural field soils were obtained from 0-30 cm pooled cores from a commercial cranberry (*Vaccinium macrocarpon* Ait.) field in Delta, BC (49.1021° N, 123.0257° W). Grassland soils were collected from randomly selected sub-pastures within two pastures at the University of Alberta Mattheis Research Ranch (50.8947° N, 111.9460° W). At the time of collection, two cores were taken within a 2.5 x 2.5 m^2^ plot in each sub-pasture for each sample, and were stored at -80 °C until analysis. At the time of analysis, multiple biological replicates from each soil type (n = 9, greenhouse; n = 10, agricultural field; n = 12, grassland) were pooled to produce single representative samples for each soil class that reflected the range of soil conditions and biological variation likely found within it. Prior to extraction, samples were sifted through a #8 mesh soil sieve to remove rocks, perlite, and plant material. All analysis was performed on subsamples of pooled soils as technical replicates to evaluate the performance of the method. A quality control sample was prepared from all soil types extracted by pooling equal volumes of each sample in a batch after extraction.

### Soil Extraction

A version of this method is freely available at DOI: https://dx.doi.org/10.17504/protocols.io.eq2lyoy6wgx9/v1. For soil extractions, 10 mL packed volume of soil from each type (greenhouse, field, or grassland) was accurately weighed. Because phytohormones are transient and thermosensitive, soils were not dried prior to extraction. Instead, soil was separately assessed for moisture content using the gravimetric method by accurately weighing 2 g of soil for each soil type and drying at 70 °C in a Kwayso dehydrator for 24 h. Moisture content was then calculated using equation 7, and soil mass was converted to dry soil equivalents in final calculations using equation 8.

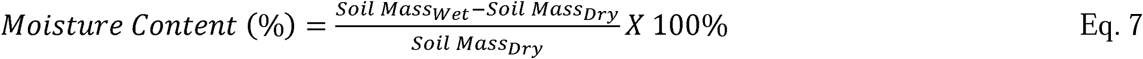

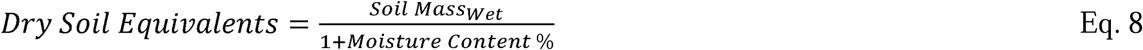

Soils were extracted by adding 20 mL of 50 % methanol (MeOH) with 0.2 % formic acid (v/v). Samples were vortexed for 5 min three times, with 5 min rest on ice, then sonicated for 10 min in a sonicator in ice water. Samples were centrifuged for 5 min at 3000 x g in an Avanti J-15 centrifuge (Beckman Coulter). The supernatant was quantitatively removed, then soils were rinsed with 5 mL of the same solvent, vortexed for 1 min, centrifuged and supernatant removed again as above. After that, 20 mL of 0.01 M NaOH was subsequently added to the soil pellet and extraction steps repeated. Supernatants were centrifuged separately for 20 min at 4000 x g, pooled, and centrifuged again to remove any retained silt. The pooled supernatants were diluted with MilliQ grade water for a final concentration of 5 % MeOH and cooled on ice prior to solid phase extraction.

Solid phase extraction was performed using Oasis HLB 3cc cartridges (Waters) containing 60 mg sorbent installed on a Supelco™ Visiprep™ SPE vacuum manifold (MilliporeSigma), with disposable liners, held at 5 mmHg. Cartridges were conditioned with 2 mL of MeOH followed by 2 mL MilliQ grade water, then samples were loaded at a flow rate of 5 mL min^-1^. Sorbent was washed with 2 mL of 5 % MeOH, then eluted twice with 500 μL 100 % MeOH. Eluted samples were loaded into Corning® Costar® Spin-X® 0.22 μm nylon spin filters (8169) and centrifuged at 16,000 x g in an Accuspin Micro17 microcentrifuge (Fisher Scientific) for 1 min to remove any remaining particulate. Filtrate was loaded into 2 mL autosampler vials for UPLC-qTOF-MS/MS analysis. An extraction method blank was prepared identically to soil samples by following all steps but without including soil. Quality control samples were analysed at the beginning, middle, and end of each analytical batch to account for signal drift during the run. As part of quality control, each batch was run with a matrix-matched standard (50 ng), typically a spiked-recovery soil sample that was used to align and verify identity of analytes in comparison to standards during processing.

### UPLC-qTOF-MS/MS Analysis

Samples were separated with an Elute Plus Ultra-high Performance Liquid Chromatography System coupled with detection via an Impact II qTOF Mass Spectrometer using a Vacuum Insulated Probe Heated Electrospray Ionization source (Bruker). Samples were separated a Bruker Intensity Solo 2 2.1 mm x 100 mm C18 column at 45 °C and using a gradient elution of 0 – 1 min 98 % A: 2 % B, 6 min 90 % A: 10 % B, 15 min 2 % A: 98 % B, 18 min 2 % A: 98 % B, 18.1 min 98 % A: 2 %B, 20 min 98 % A: 2 % B at a flow rate of 0.3 mL min^-1^. The solvents were 0.1 % formic acid in water (A) and 0.1 % formic acid in acetonitrile (B), and samples were injected at a volume of 20 μL. Between each sample, the needle was washed with 1.5 mL of solvent A and 1.5 mL of 90 % acetonitrile to prevent carryover. Between 0.05 min and 0.3 min, 10 μL sodium formate (containing isopropanol, 0.01 M sodium hydroxide, and formic acid at a ratio of 50:50:0.2) was injected as an external mass calibrant. Samples were separately run in positive and negative mode on full MS and autoMSMS scanning. The source was set to 2.0 bar (nebulizer gas); 8.0 L min^-1^ (dry gas flow); 4.0 L min^-1^ (probe gas flow); 230 °C (source temperature); 400 °C (sheath gas/probe gas temperature), 4500 V (capillary voltage), at 20 – 700 m/z scan range using a spectral rate of 12.00 Hz. For autoMSMS mode, a 5, 10, and 15 eV collision energy series was utilized along with an ion energy of 5 eV, prepulse storage time of 3 μs, transfer time of 45 μs, radio frequencies of 450 Vpp (collision), 200 Vpp (funnel 1 and 2), and 50 Vpp (hexapole), and a cycle time of 0.3 s with an absolute threshold of 400 counts per 1000 sum. A scheduled precursor list was used to define precursors for MS/MS, with the number of precursors set to 3 and MS/MS spectra acquired in empty intervals (Table 2). MS/MS spectra was set to dynamic with a target intensity of 20000 counts and a spectral acquisition range of 16.00 – 30.00 Hz.

**Table 2:** Scheduled precursor lists for acquisition of MS/MS spectra in positive and negative ionization modes.

| Analyte<sup>1</sup> | RT<sup>2</sup> (min) | Positive Ionization |  | Negative Ionization |  |
| --- | --- | --- | --- | --- | --- |
|  |  | RT Tolerance | Mass Range | RT Tolerance | Mass Range |
| ABA | 10.10 | 0.05 | 265.1240 - 265.1640 | 0.05 | 263.1084 - 263.1484 |
|  |  | 0.05 | 247.1128 - 247.1528 | 0.05 | 527.2298 - 527.2698 |
| IAA | 9.55 | 0.05 | 176.0511 - 176.0911 | 0.10 | 153.3600 - 153.4000 |
|  |  | 0.05 | 130.0465 - 130.0865 | 0.10 | 174.0400 - 174.0800 |
| IBA | 10.60 | 0.05 | 186.0718 - 186.1118 | 0.15 | 202.0668 - 202.1068 |
|  |  | 0.05 | 204.0824 - 204.1224 | 0.15 | 405.1499 - 405.1899 |
| JA | 10.80 | 0.05 | 211.1134 - 211.1534 | 0.05 | 209.0978 - 209.1378 |
|  |  | 0.05 | 193.1015 - 193.1415 | 0.05 | 165.1053 - 165.1453 |
| KAR1 | 9.00 | 0.05 | 151.0190 - 151.0590 | NA | NA |
| KAR2 | 5.90 | 0.10 | 137.0033 - 137.0433 | NA | NA |
| KAR3 | 10.20 | 0.05 | 165.0346 - 165.0746 | NA | NA |
| KAR4 | 10.00 | 0.05 | 165.0346 - 165.0746 | NA | NA |
| Mel | 9.45 | 0.05 | 233.1090 - 233.1490 | 0.10 | 231.0934 - 231.1334 |
|  |  | NA | NA | 0.10 | 216.0631 - 216.1031 |
| BA | 8.65 | 0.05 | 226.0892 - 226.1292 | 0.10 | 224.0736 - 224.1136 |
|  |  | NA | NA | 0.10 | 133.0168 - 133.0568 |
| 2IP | 8.45 | 0.05 | 204.1049 - 204.1449 | 0.10 | 202.0893 - 202.1293 |
|  |  | NA | NA | 0.10 | 158.0242 - 158.0642 |
| Trp | 5.20 | 0.10 | 205.0777 - 205.1177 | 0.15 | 203.0621 - 203.1021 |
|  |  | NA | NA | 0.15 | 142.0449 - 142.0849 |
| Zea | 4.85 | 0.10 | 220.0998 - 220.1398 | 0.15 | 218.0842 - 218.1242 |
| GA3 | 8.20 | NA | NA | 0.15 | 345.1138 - 345.1538 |
|  |  | NA | NA | 0.15 | 283.1067 - 283.1467 |
| SA | 9.50 | NA | NA | 0.15 | 137.0039 - 137.0439 |
|  |  | NA | NA | 0.15 | 93.0131 - 93.0531 |

<sup>1</sup>ABA: Absciscic acid; IAA: indole-3-acetic acid; IBA: indole-3-butyric acid; JA: jasmonic acid; KAR1 – 4: karrikins 1-4; Mel: melatonin; BA: N6-benzyladenine; 2IP: N6-isopentenyladenine; Trp: tryptophan; Zea: zeatin; GA3: gibberellic acid A3; SA: salicylic acid.
<sup>2</sup>RT: Retention time;

For quantification, a standard curve typically consisting of 5 points covering the range of 0 – 50 ng mL^-1^ for all spiked analytes was selected based on the established linear range. Raw data were quantified using Bruker TASQ 2024 software for targeted analysis. The Peak Fit algorithm was used for finding peaks, with height for calculating ion ratios and the Expert algorithm for calculating signal to noise parameters, using default settings recommended for the Impact II. During processing, signals were smoothed and denoised, then compared to an in-house target list for identification (Table S2). Identification was performed using principal ion for ion quality. Determination filters were set to include most mandatory ions and those closest to retention times, with a narrow tolerance of 0.25 min and wide tolerance of 0.60 min. All ions had a sensitivity of 99%, minimum peak valley of 4%, ppm window of 3.00 (narrow) and 7.00 (wide), and a smoothing width of 1 s. For [M+H+1]^+^ or [M-H+1]^-^ ions, listed ion ratios were set using [M+H]^+^ or [M-H]^-^ ions as reference and with a tolerance of 30 %. Signals were quantified using peak area, with area thresholds listed in Table S2. Quantification was typically performed using the principle ion, [M+H+1]^+^ or [M-H+1]^-^ ions, and the second most abundant ion based on FullScan MS spectra. As part of quality control, peak identities, peak picking and quantification were evaluated for consistency and accuracy through manual inspection and comparison to matrix-matched standard.

For untargeted metabolomics, samples were processed through Metaboscape 2025 (Bruker). Peaks were detected using intensity as the feature signal, an intensity threshold of 1E4, a minimum peak length of 10 spectra, and with recursive feature extraction enabled using the same minimum peak length. For deconvolution, an EIC correlation of 0.8 was used. Finally, samples were calibrated using an internal calibration signal between 0-0.3 min of Na formate clusters. After the feature table was generated, blank subtraction was performed using the extraction blank and a solvent blank consisting of 50 % MeOH, then false peaks (artifacts) were excluded following manual inspection.

Peaks were annotated by matching to an in-house hormonomics target list based on the Hormonomics Database and the following positive and negative spectral libraries: Plant Metabolites, Bruker Sumner MetaboBASE Plant Library (confirmed standards, putative ID, and NMR confirmed), MS/MS Vaniya Fiehn Natural Products Library (2020), MS/MS MetaboBASE, MS/MS MassBank, MS/MS Global Natural Products (GNPS), MS/MS Bioinformatics & Molecular Design Research Center Mass Spectral Library (Natural Products), MS/MS Pathogen Box, BioMS/MS PlaSMA, MS/MS Public (Exp,Exp VS17 and Exp Bio in silico), MS/MS PlaSMA, MS/MS Respect, MS/MS Critical Assessment of Small Molecule Identification (2016), and MS/MS MassBank EU (Erland et al., 2020; Tsugawa et al., 2019, 2015). Annotated features were exported to Metaboanalyst 6.0 for further analysis (Pang et al., 2024). The statistical analysis module (one factor) was used on the annotated features list after peak area had been normalized to moisture-corrected dry weight. During data import, the dataset was normalized to sum, square-root transformed to reduce the presence of outliers, then scaled using Pareto scaling. A principal component analysis was performed, as well as Analysis of Variance using a false discovery rate cutoff of 0.05 in the related functionalities. A heat map was generated using Euclidean distance and Ward’s D on the significant features identified through ANOVA.

### Statistical Analysis

As described above, all experiments were performed using composite soil samples representing each soil class. Composite soil samples were prepared by pooling biological replicates representative of research activities taking place at each site (n = 9, greenhouse; n = 10, agricultural field; n = 12, grassland) to capture the range of variation likely to occur within each soil class. Youden analysis (n = 3), matrix effects (n = 4), and spiked recovery (n = 4, greenhouse and agricultural field; n = 5, grassland), as well as targeted and untargeted analysis were all performed using these composite soil samples to showcase variation in the method. Method validation was completed using triplicate injections, except for LOD/LOQ calculations which were performed with replicate blank injections (n = 8, negative ionization; n = 9, positive ionization). Processing of quantitative data was performed with TASQ 2025 (Bruker), and untargeted metabolomics performed with Metaboscape 2025 (Bruker) and Metaboanalyst 6.0 (Pang et al., 2024). All other graphical and statistical analyses were performed using GraphPad Prism 10.

## Results

### Method Optimization and Validation

Prior to performing a Youden analysis, optimization showed that a sequential extraction utilizing both acidic and basic solvents recovered more analytes than using acidic or basic extractions alone, and a packed soil volume of 10 mL was needed to establish a baseline signal in un-spiked soil samples. The smaller sorbent mass was selected to reduce the volume of eluent required to fully remove spiked standards from the sorbent during SPE. Youden analysis revealed the most influential factors investigated were acid choice, extraction order, and base concentration, but this was limited to a few analytes (Figure 2). Acid choice was not a main effect on recovery for most analytes but had a strong effect on the recovery of KAR2, which was favored by acetic acid extraction. A higher concentration of NaOH was more effective at recovering ABA, JA, KAR3, and SA up to 215%, and the order of extraction was more important for ABA, KAR1, KAR2, SA, GA3 and JA, with up to a 4.3-fold increase in analyte measured when extracted with acid first. The opposite trend was recorded for Zea, which was not detected unless basic conditions were applied first. Trp also benefited from base-first extraction, which increased measured analyte by 131% compared to acid-first extractions. The addition of salt to the basic extraction solvent had very little effect on any analyte except for KAR2, which was only detected when salt was not present. Prewetting likewise had little impact on recovery. Extracted ABA, JA, and Trp increased by 55, 59, and 88% respectively with shorter sonication time, while longer sonication times considerably increased extracted karrikins by up to 369%.

**Figure 2.**
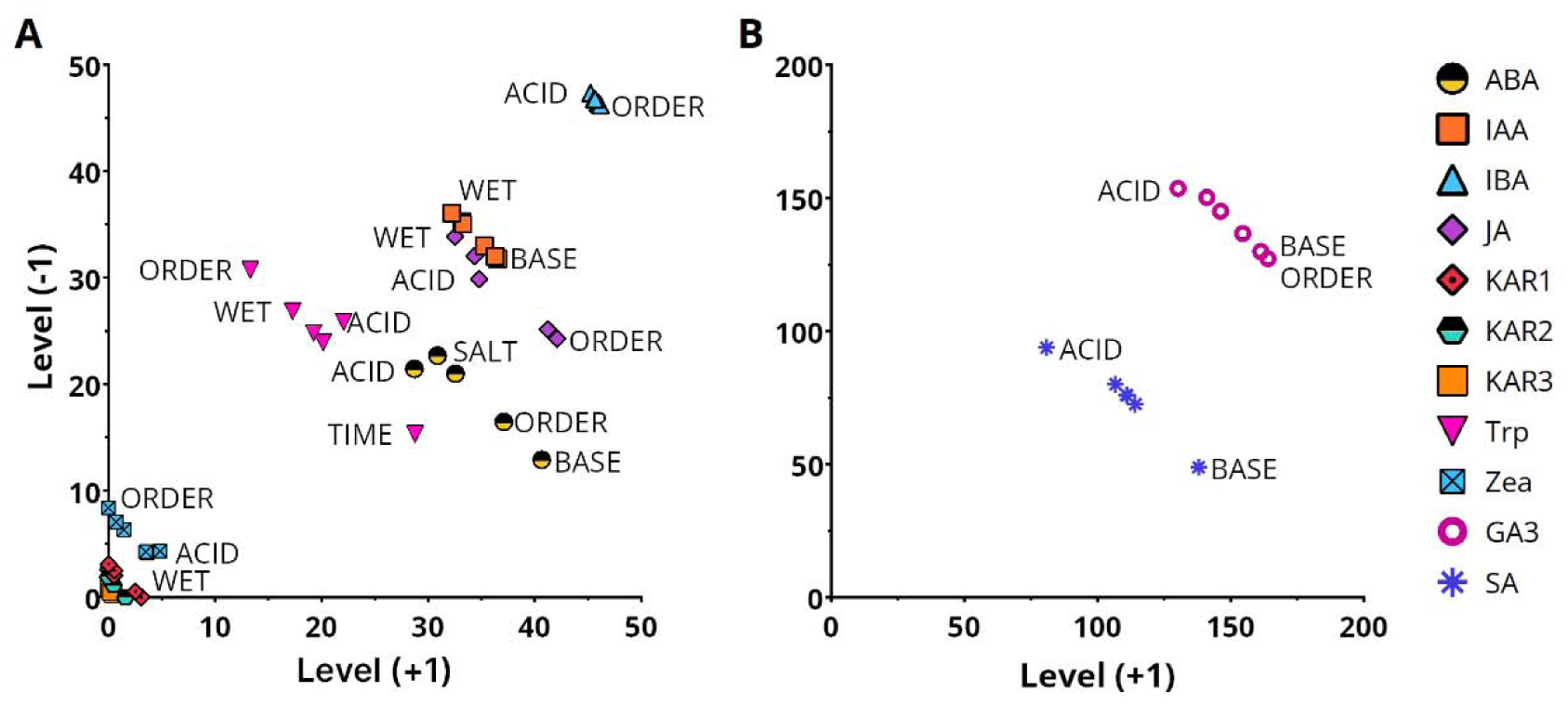
Youden analysis of 6 main factors and their effect on mean measured quantity (pg/g) of 9 phytohormones recovered from spiked soil extractions (n=3) using clay-rich agricultural field soil. **A**. Lower abundance hormones and **B**, higher abundance hormones, letters reflect main factor effects. For clarity, only the most influential factors are labeled. Alt-text: Two scatter-plots depicting the output of a Youden analysis for 11 phytohormones. The x and y axes indicate the level a factor was tested at, with the lowest level on the y-axis and the highest level on the x-axis.

From this analysis, the method parameters were selected to maximize recovery across all analytes: extraction proceeded with 50% MeOH with 0.2% formic acid followed by 0.01N NaOH with an extraction cycle of 3 x 5 min vortexing/5 min rest on ice, 10 min sonication with no additional salt or prewetting steps required. Four percent acetic acid or 0.1N NaOH can be flexibly substituted when karrikin and general hormones such as ABA or SA enrichment is desirable, respectively.

Across positive and negative ionization modes, all analytes were detectable at a high degree of linearity, with most analytes at an R^2^ value of 0.990 or higher (Table 3) across a range of 3 orders of magnitude. KAR3, KAR4 and Mel had lower values of linearity between 0.981 – 0.989, which may be attributable to co-elution with other analytes (KAR3, KAR4, Figure 3). This is also reflected in inter-day repeatability measures, with most analytes below 8.9 % RSD but KAR2, Trp, and Zea showing higher inter-day variability up to 34.80 % (Trp). Analytes were detectable at a range of 0.01 – 15,000 pg on-column. The highest degree of sensitivity was shown for ABA, with an LOQ of 1.4 pg on-column (negative ionization), while the analyte with the least sensitivity was salicylic acid (negative ionization, 167.3 pg on-column). Accuracy of the extraction method varied by soil type (Table 4). Matrix-adjusted recovery rates in grassland soils ranged from 1.0 – 154.0%, in greenhouse soils from 5.5 – 99.8%, and in agricultural field soils from 0.1 – 306.6%. Although the range of recovery rates was similar across soil types, recovery rates were generally higher in grassland soils and lowest in agricultural field soils. ABA in agricultural field soils and GA3 in grassland soils were over-recovered when accounting for ion suppression effects (306.6 ± 64.4 and 154.0 ± 15.0 %), while Mel, BA, 2IP, Trp, and Zea were poorly recovered in comparison.

**Figure 3.**
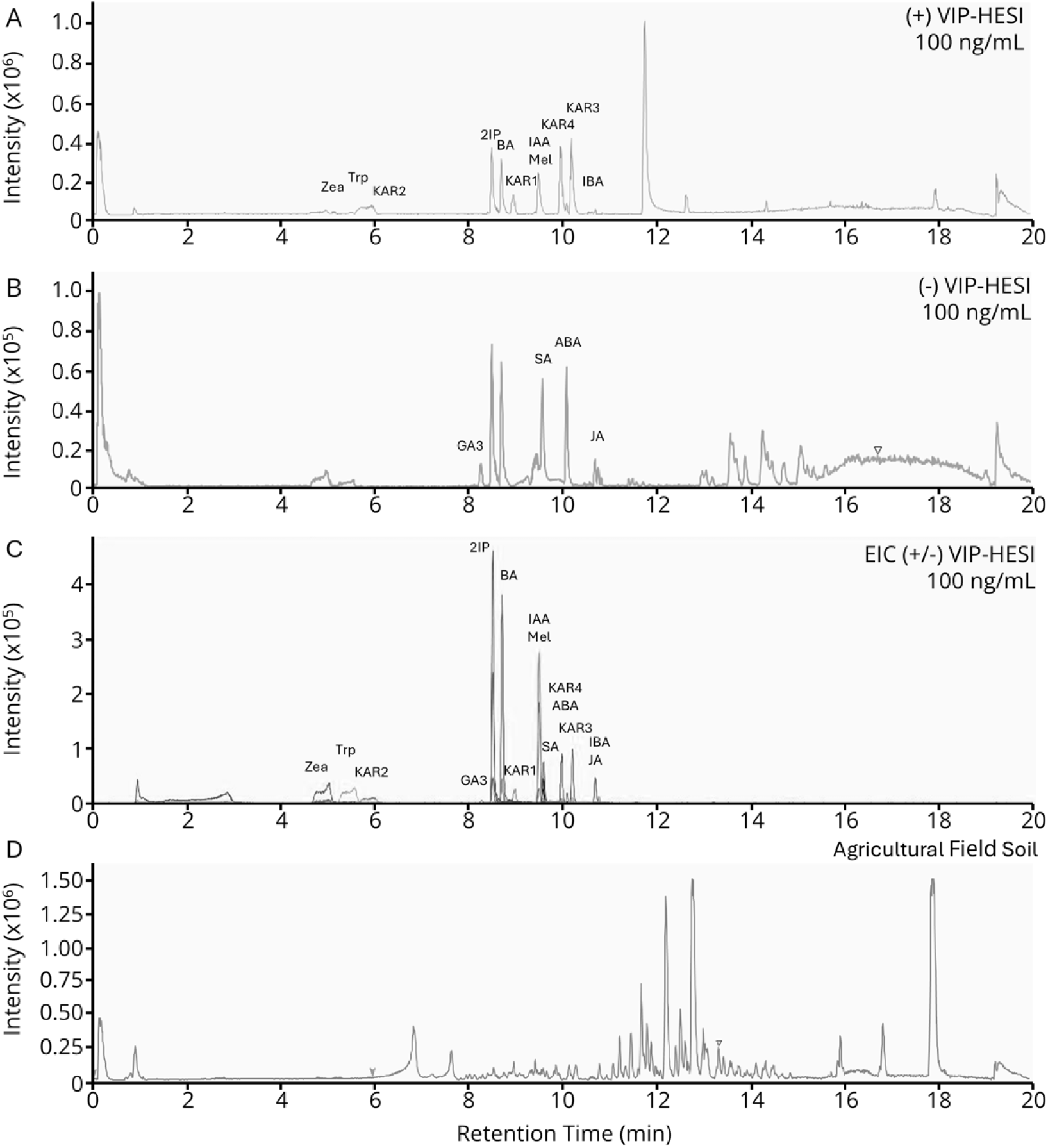
Typical base peak chromatograms for standard mix containing ABA, IAA, IBA, JA, KAR1, KAR2, KAR3, KAR4, Mel, BA, 2IP, 5HT, Trp, Zea, SA and GA3 on positive ionization (**A**) and negative ionization modes (**B**). Extracted ion chromatograms from positive and negative ionization used for quantitation are shown in **C**. A typical base peak chromatogram of an extraction for agricultural field soil in positive ionization is shown in **D**. ABA: Abscisic acid; IAA: indole-3-acetic acid; IBA: indole-3-butyric acid; JA: jasmonic acid; KAR1 – 4: karrikins 1-4; Mel: melatonin; BA: N6-benzyladenine; 2IP: N6-isopentenyladenine; Trp: tryptophan; Zea: zeatin; GA3: gibberellic acid 3; SA: salicylic acid. Alt-text: A series of LC-MS chromatograms showing the intensity of signal for positive and negative ionization, and a stacked extracted ion chromatogram for 15 analytes plotted against retention time in minutes. A separate chromatogram is shown for a typical soil extraction.

**Table 3:** Parameters of method validation including limit of detection (LOD), limit of quantitation (LOQ), range, linearity and inter-and intra-day precision. LOD/LOQ was calculated from replicate blank injections (positive ionization, n = 9; negative ionization, n = 8). Parameters were calculated from three replicate injections of a hormone standard containing a mix of all analytes over a 5-point (inter-day %RSD, n = 3 per day injected) or 12-point (all other parameters) standard curve.

| Analyte<sup>1</sup> | Ionization Mode | RT<sup>2</sup><br>(min) | LOD | LOQ | Range | Slope | y-intercept | R2 | %RSD |  |
| --- | --- | --- | --- | --- | --- | --- | --- | --- | --- | --- |
|  |  |  |  |  |  |  |  |  | Inter-day | Intra-day |
| ABA | NEG (-) | 10.10 | 0.005 | 1.41 | 1.4 - 5000 | 2066 | 11444 | 0.996 | 7.1 | 2.4 ± 1.7 |
| IAA | POS (+) | 9.55 | 4.7 | 15.5 | 15.5 - 10000 | 2831 | 14937 | 0.996 | 4.5 | 2.5 ± 1.5 |
| IBA | POS (+) | 10.60 | 5.3 | 17.5 | 17.5 -10000 | 2230 | 14842 | 0.996 | 7.0 | 2.7 ± 0.5 |
| JA | NEG (-) | 10.80 | 31.68 | 105.61 | 105.6 - 15000 | 345.6 | 1077 | 0.999 | 6.4 | 2.7 ± 1.3 |
| KAR1 | POS (+) | 9.00 | 4.8 | 16 | 16 - 1000 | 8538.7 | 16295 | 0.994 | 3.4 | 3.2 ± 1.4 |
| KAR2 | POS (+) | 5.90 | 0.9 | 2.9 | 2.9 - 1500 | 15953 | 19414 | 0.994 | 27.7 | 9.6 ± 3.0 |
| KAR3 | POS (+) | 10.20 | 1.4 | 4.8 | 4.8- 500 | 24524 | 36591 | 0.989 | 5.0 | 2.0 ± 1.2 |
| KAR4 | POS (+) | 10.00 | 1.3 | 4.2 | 4.2- 1000 | 21654 | 31789 | 0.988 | 8.2 | 4.5 ± 1.7 |
| Mel | POS (+) | 9.45 | 0.8 | 2.6 | 2.6 - 5000 | 11495 | 121042 | 0.981 | 5.7 | 1.3 ± 1.3 |
| BA | POS (+) | 8.65 | 39 | 130 | 130 - 5000 | 11725 | 66419 | 0.994 | 8.9 | 4.6 ± 3.4 |
| 2IP | POS (+) | 8.45 | 0.9 | 2.9 | 2.9 - 5000 | 18352 | 119717 | 0.993 | 7.8 | 6.2 ± 2.9 |
| Trp | POS (+) | 5.20 | 2.4 | 7.9 | 7.9 - 10000 | 5280 | -29266 | 0.998 | 34.8 | 7.0 ± 4.5 |
| Zea | POS (+) | 4.85 | 8.7 | 29 | 29 - 2000 | 4665 | 60461 | 0.991 | 18.9 | 4.3 ± 2.5 |
| GA3 | NEG (-) | 8.20 | 2.93 | 9.76 | 9.8 - 10000 | 647.5 | 2050 | 0.998 | 17.2 | 9.3 ± 7.4 |
| SA | NEG (-) | 9.50 | 50.19 | 167.3 | 167.3 - 10000 | 2067 | 53203 | 0.990 | 9.8 | 2.3 ± 1.5 |

<sup>1</sup>ABA: Absciscic acid; IAA: indole-3-acetic acid; IBA: indole-3-butyric acid; JA: jasmonic acid; KAR1 – 4: karrikins 1-4; Mel: melatonin; BA: N6-benzyladenine; 2IP: N6-isopentenyladenine; Trp: tryptophan; Zea: zeatin; GA3: gibberellic acid A3; SA: salicylic acid
<sup>2</sup>RT: Retention time

**Table 4:** Matrix-adjusted recovery rates for 15 phytohormones in three soil types prepared by pooling biological replicates (n = 9, greenhouse; n = 10, agricultural field; n = 12, grassland) from each research site to create a representative sample. Matrix effects were determined by evaluating post-extraction spike recovery samples (n = 4) spiked with 50 ng/mL of listed analytes for each soil class. Matrix-adjusted recovery rates were determined from analysis of spike recovery samples (greenhouse, agricultural field, n = 4; grassland, n = 5) spiked with 50 ng of each analyte.

| Analyte<sup>1</sup> | Matrix - Adjusted Recovery Rate (%) |  |  | Matrix Effect (%) |  |  |
| --- | --- | --- | --- | --- | --- | --- |
|  | Grassland | Greenhouse | Ag. Field | Grassland | Greenhouse | Ag. Field |
| ABA | 99.7 ± 4.7 | 81.2 ± 6.8 | 306.6 ± 64.4 | 49 ± 5 | 27 ± 4 | 3 ± 7 |
| IAA | 56.0 ± 6.8 | 66.1 ± 5.0 | 24.0 ± 1.4 | 44 ± 2 | 22 ± 5 | 17 ± 4 |
| IBA | 57.2 ± 8.1 | 36.4 ± 1.2 | 1.1 ± 2.2 | 63 ± 12 | 34 ± 1 | 29 ± 2 |
| JA | 100.4 ± 22.4 | 99.8 ± 18.9 | 132.0 ± 21.0 | 46 ± 7 | 17 ± 8 | 5 ± 7 |
| KAR1 | 97.1 ± 6.7 | 64.0 ± 3.7 | 36.4 ± 0.5 | 30 ± 3 | 19 ± 2 | 17 ± 1 |
| KAR2 | 57.6 ± 1.6 | 46.1 ± 6.5 | 23.4 ± 7.0 | 24 ± 10 | 17 ± 1 | 17 ± 12 |
| KAR3 | 77.8 ± 13.6 | 35.0 ± 3.4 | 0.0 ± 0.0 | 36 ± 1 | 16 ± 2 | 13 ± 0 |
| KAR4 | 81.3 ± 19.7 | 58.1 ± 5.8 | 12.6 ± 1.7 | 30 ± 1 | 13 ± 1 | 11 ± 2 |
| Mel | 41.5 ± 4.7 | 22.2 ± 11.2 | 0.0 ± 0.0 | 38 ± 5 | 25 ± 4 | 17 ± 7 |
| BA | 1.3 ± 0.4 | 5.5 ± 0.5 | 0.1 ± 0.0 | 120 ± 6 | 85 ± 8 | 96 ± 5 |
| 2IP | 1.0 ± 0.1 | 12.1 ± 0.4 | 2.3 ± 0.1 | 91 ± 4 | 69 ± 12 | 76 ± 10 |
| Trp | 12.9 ± 1.8 | 14.1 ± 1.3 | 8.2 ± 0.8 | 40 ± 6 | 48 ± 11 | 65 ± 17 |
| Zea | 8.5 ± 0.1 | 21.2 ± 5.2 | 8.5 ± 0.4 | 46 ± 10 | 50 ± 9 | 57 ± 18 |
| GA3 | 154.0 ± 15.0 | 87.7 ± 24.9 | 59.8 ± 15.0 | 54 ± 8 | 41 ± 8 | 40 ± 4 |
| SA | 74.2 ± 6.3 | 83.6 ± 13.1 | 60.0 ± 12.4 | 43 ± 5 | 34 ± 7 | 25 ± 10 |

<sup>1</sup> ABA: Absciscic acid; IAA: indole-3-acetic acid; IBA: indole-3-butyric acid; JA: jasmonic acid; KAR1 – 4: karrikins 1-4; Mel: melatonin; BA: N6-benzyladenine; 2IP: N6-isopentenyladenine; Trp: tryptophan; Zea: zeatin; GA3: gibberellic acid A3; SA: salicylic acid.

### Quantitative Analysis of Soils

Subsamples of pooled grassland, greenhouse and agricultural field soil were analysed for each analyte to establish endogenous phytohormone levels. Soil type and texture varied strongly among soils (Figure 4A). This corresponded to differences in the presence and abundance of endogenous phytohormones for each soil type. Analytes are represented as a proportion of total phytohormones quantified using the targeted method (Figure 4B). In grassland soil, the phytohormone profile is dominated by SA and IAA at 34.3 and 6.8 pg/g moisture-corrected dry weight respectively, with trace quantities of ABA (1.5 pg/g) and JA (< LOQ). Greenhouse soil had the greatest diversity of phytohormones, with ABA, IAA, Trp, Mel, JA, Zea and SA detected. The most abundant phytohormone was SA (586.6 pg/g), followed by IAA (283.9 pg/g), ABA (163.4 pg/g), Trp (131.7 pg/g), and Mel (74.1 pg/g). JA and Zea were present only at trace quantities, and below the limit of quantitation. Agricultural field soil also contained high amounts of SA (409.2 pg/g), as well as trace amounts of IAA, JA, and Trp that were detected but could not be quantified.

**Figure 4.**
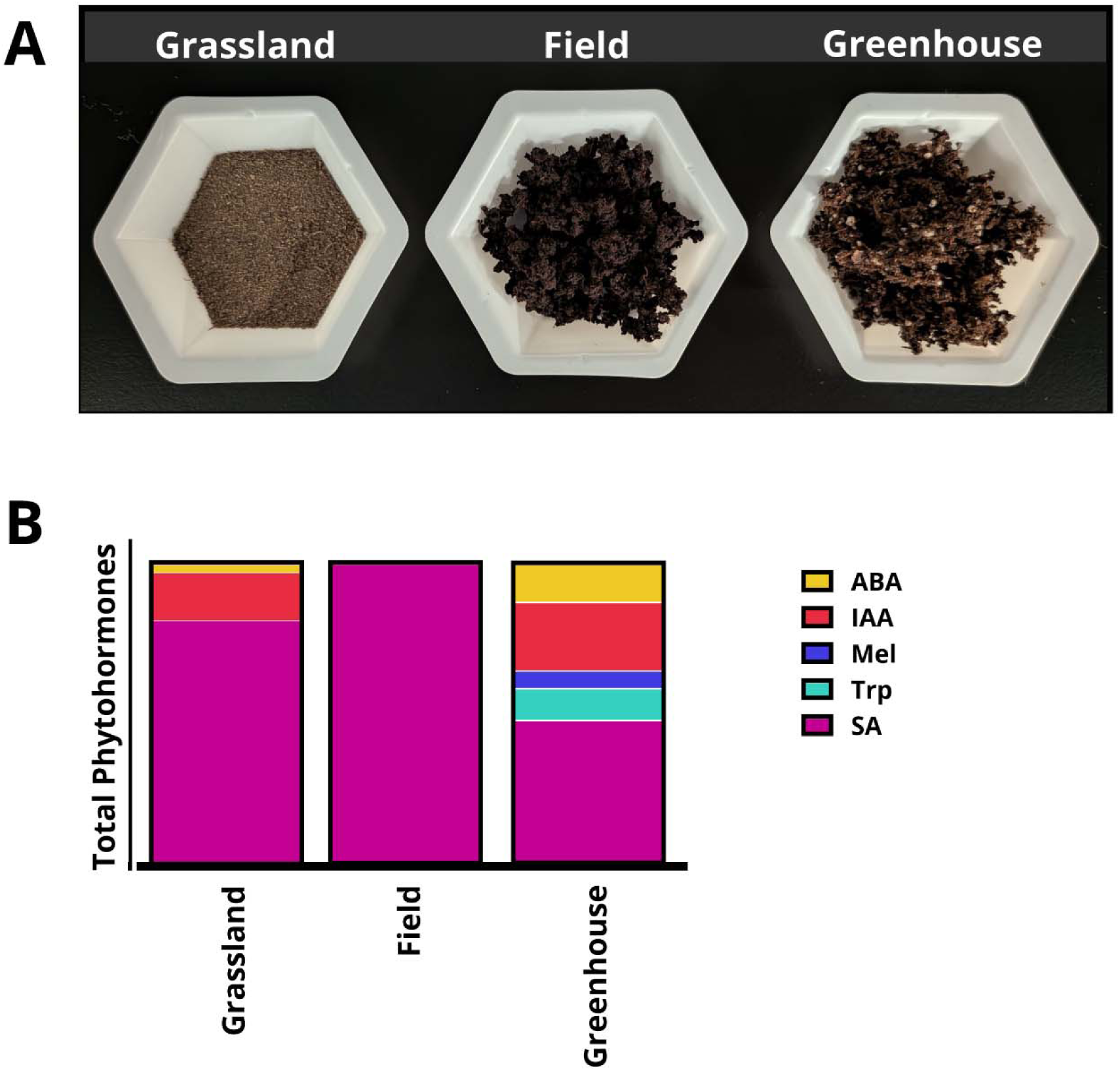
**A.** Representative samples of soils from grassland, agricultural field and greenhouse soils. **B.** Proportional representation of endogenous phytohormone abundance in soil extractions (n = 3) for each soil type (n = 3). ABA: Abscisic acid; IAA: indole-3-acetic acid; Mel: melatonin; Trp: tryptophan; SA: salicylic acid. Alt-text: A photographic image showing soil texture for three soils is shown in panel A. Grassland soil has a fine, sandy texture, while field soil is dense and clay-rich and greenhouse soil is light and peat-rich. In panel B, three stacked bars show abundance of phytohormones for the same three soils, with color-coding to identify analyte and bar size indicating proportional abundance compared to total phytohormones.

### Untargeted Analysis of Soils

Using the same subsamples we used for targeted analysis, we further analysed them using an untargeted workflow. After processing, one hundred and two mass features were annotated using an in-house target list for hormones and their derivatives, as well as to available public libraries (Table S3). A principal component analysis was conducted followed by PERMANOVA (F = 1078.5, R^2^ = 0.9972) to establish the consistency of the extraction method on annotated mass features. Samples from each soil type separated with like samples (Figure 5A). Principal component 1 accounted for 50.1 % of variation, and PC2 34.4 %. Separation along PC1 was strongly driven by several agriculturally relevant man-made compounds (methoxyfenozide, an insecticide, metalaxyl metabolites and trifloxystrobin (fungicides), and chlorantraniliprole (an insecticide), along with coumaric and ferulic acids and precursors for JA, such as 12-oxo-phytodienoic acid that were abundant in agricultural field soils. Separation along PC2 was driven by the same anthropogenic compounds, but additionally by several flavonoids such as 1-(3,4-dihydroxyphenyl)-7-(4- hydroxyphenyl)heptan-3-one, and isobavachalcone, a compound produced by members of the genus *Glycyrhizia.* An ANOVA revealed 78 annotated mass features that differed between soil types (Table S4). These were visualized as a heat map using Euclidean distance and Ward’s D agglomerative clustering, with subsamples shown as group averages (Figure 5B). Clustering reveals three groups of annotated features that provide a unique profile for each soil. Grassland soils show differential abundance of 9,10-dihydrojasmonate, DL-β-hydroxybutyric acids, and linoleic acid, in addition to isobavachalcone (Figure 5C). Agricultural field soils are enriched in cytokinin derivatives such as topolin and topolin ribosides, tryptophan-derived metabolites like kynurenine, and an abundance of phenylpropanoids including fraxedin, fraxin, ferulic acids, and the iridoid monoterpene asperuloside commonly found in the genus *Galium*. Greenhouse soils show a differential abundance of JA precursors and derivatives such as 9,10-dihydrojasmonic acid, in addition to a selection of agricultural pesticides and fungicides.

**Figure 5.**
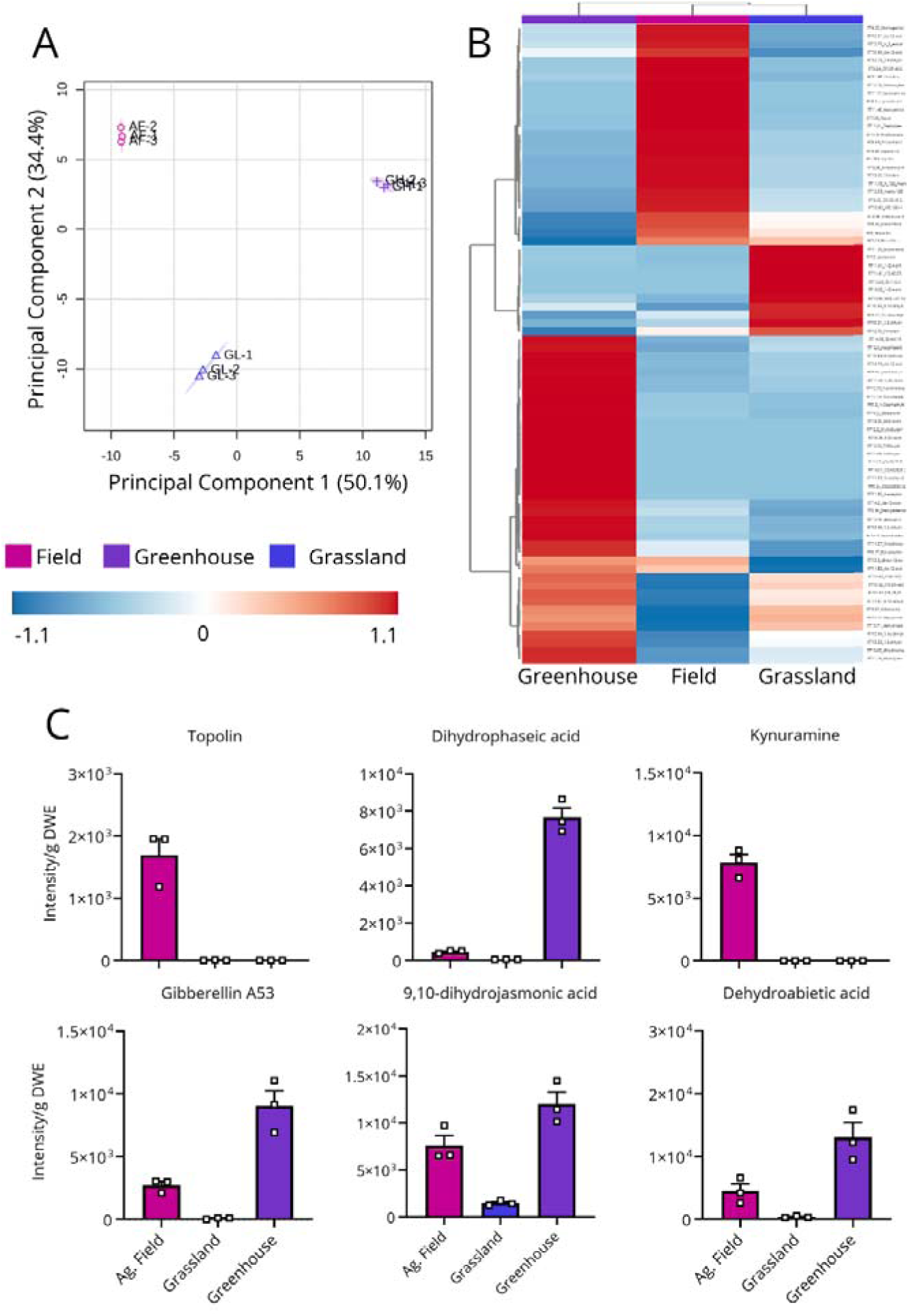
Untargeted hormonomic analysis of replicate soil extractions (n =3) for composite samples of each soil class prepared from pooled biological replicates taken from each research site (n = 9, greenhouse, n = 10, agricultural field, n = 12, grassland) **A**. Principal component analysis shows soil samples separate into groups by soil type with low variance. **B.** A heat map prepared with Euclidean distance and Ward’s D agglomerative clustering from 78 annotated features identified as significant following Analysis of Variance with a false discovery cutoff of 0.05 shows soil types form into distinct clusters. **C.** Representative features of phytohormones annotated using in-house target list that are differentially abundant across soil types are shown. Error bars indicate standard error. Alt-text: A three-panel figure showing graphs related to untargeted metabolomic analysis. Panel A shows the results of a principal component analysis for principal component 1 on the x-axis and 2 on the y-axis. Panel B depicts a heat map of annotated mass features identified in three soil types, and resolved into three clusters by soil. The heat map is visualized with a two-color gradient where blue indicates differentially lower and red indicates differentially increased abundance of each feature. Panel C shows bar graphs of six phytohormones annotated by target list. Each bar graph depicts the abundance of a phytohormone on the y-axis in units of peak intensity per gram dry weight equivalent of soil extracted, and soil type on the x-axis.

## Discussion

Soils are increasingly the new frontier of plant research, with soil metabolomics being used as a means of investigation of soil health and ecosystem processes, soil microbiome composition, markers of ecotoxicity and more (Bhattacharjya et al., 2024). Phytohormones are transient, small molecules which can help plants to perceive and respond to the world around them making them an important component and likely determinant of plant health and resilience. By expanding our understanding of phytohormone abundance and diversity to soils, we open new opportunities to understand the broader role they may play in the soil system, and for future evaluations of root and soil health. Unlocking the means to explore these important aspects of environmental-level hormone interactions will enable us to explore these concepts and how they impact stress resilience and the provisioning of ecosystem goods and services.

Both soil type and analyte influenced our extraction method. This is consistent with other methods that also see an influence of soil type on parameters such as recovery (McManus et al., 2019; Wan Mohd Zamri et al., 2021). Although overall recovery rates were high, extraction was more successful on grassland and greenhouse soils than agricultural field soils. This is likely due to the abundance of fine particulate and clay content in these soil types, which increase cation exchange capacity and causes analytes to bind more tightly to soil particles with a high surface area (Lee et al., 2025). Greenhouse soils also had lower recovery of most analytes compared to grassland soils, and this is likely due to the high degree of organic material present in peat-based soils, which contain humic and fulvic acids that are known to interfere with the extraction of small molecules (Ibáñez et al., 1998; Qian and Hettich, 2017). There was also an increase in the influence of matrix effects causing ion suppression during analysis that was dependent on soil type and analyte. Matrix effects were lowest in grassland soils and highest in agricultural field soils. This may also be attributable to high cation exchange capacity and the release of interfering metabolites. Solid phase extraction removed interferences by salts, but other metabolites like lipids can also cause significant ion suppression by interfering with droplet formation during electrospray ionization, for example, and are often co-extracted along with desired analytes during extractions (Dąbrowska et al., 2003). The use of matrix-matched standard curves may help to overcome these effects, and have been successfully used to limit the consequences of matrix effects in other analyses of soil extraction (McManus et al., 2019). We used a single matrix-matched standard as part of our quality control process, where it enabled simultaneous adjustment of analyte concentration as a spiked-recovery standard and allowed for more accurate peak-picking during processing by showcasing matrix effects on peak shape and retention time. Because of these differences in recovery rates dependant on soil type, we recommend extending this concept to the inclusion of a composite matrix-matched standard prepared from all soils of a similar type to act as a combined recovery and quality control sample during analysis.

Differences in analyte recovery between analytes is due to a number of factors. Previous work has established that hormones are sensitive to a variety of external influences in active biological and environmental systems, including exposure to light, increased temperature, and oxygen that can significantly reduce recovery of the untransformed analyte (Erland et al., 2016). For example, although melatonin on its own is relatively stable, temperature during extraction affects recovery, with increased recovery at lower temperature. Extraction time also contributes to this, as lengthy extraction steps allow time for chemical transformation prior to capture and analysis (Pan et al., 2008). Melatonin and other Trp derivatives are highly reactive antioxidants, and are extremely susceptible to not only redox reactions with coextracted metal ions, but also reactions with other desirable analytes as reactive intermediates, reducing recovery for both (Mukherjee and Ramakrishna, 2018). Both Mel, and Trp had lower recovery rates in our study that may be partially explained by this phenomenon. To counter these possible influences, we recommend performing the extraction consecutively on a single day in smaller sample batches, with all steps chilled at 4 °C and protected from light where possible. For particularly silt or particulate-rich extractions, we also recommend pre-filtering extractions prior to performing SPE to reduce interference of heavy particulate on recovery efficiency. Finally, we also noted that recovery was lower for cytokinins including Zea, 2IP and BA, which may be extracted better under basic conditions.

We used our targeted and untargeted method to evaluate the endogenous phytohormone profile of agricultural field, grassland, and greenhouse soils. All three soils contained high amounts of salicylic acid. Grassland soils had, in addition to salicylic acid, ABA and detectable but unquantified levels of JA, which are stress- and defense-related phytohormones that may indicate an ongoing stress response. For example, salicylic acid can change root architecture to influence water uptake by stimulating growth, especially under drought stress (Harris, 2015), while exogenous application of jasmonates have been shown to increase resilience of plant to salt stress in several crop species (Delgado et al., 2021; Harris, 2015). This may also indicate biotic challenges, since JA and its derivatives are activated by plants in response to a range of different pathogens, including bacteria, fungi, and oomycetes but also in response to endophytic colonization (Chen et al., 2020; Delgado et al., 2021; Thaler et al., 2004). These soils also had phytochemicals in the untargeted analysis which may provide indication of plant community composition. For example, in agricultural field soils we found a high abundance of asperuloside, which is found in *Galium sp*. *Galium verum* is a naturalized herbaceous species likely introduced to Canada from Europe, and is found in high abundance in Alberta and British Columbia where it is likely to become an invasive species (Mersereau and DiTommaso, 2003). In grassland soils, metabolites found in species including *Glycyrhizia* were noted, members of which can be found as a native plant throughout Alberta. Both greenhouse and agricultural field soils contain metabolites that link to their use as two commercial crop settings, including common fungicides and insecticides, but these soils are also highly dynamic, enriched in cytokinins, jasmonates, and plant growth regulators indicative of active growth. The abundance of salicylic acids and jasmonates indicates ongoing abiotic and biotic stress responses in these samples, demonstrating the potential for soil phytohormone analysis as a mean to characterize plant and plant community health.

Collectively, the paired targeted and untargeted analyses presented here are complimentary, providing a more comprehensive understanding of phytohormones and other active metabolites in these soil types that relate to their environment. Our targeted analyses allow for direct quantitative comparison between soils, while untargeted analysis provide hypothesis-generating, fit-for-purpose relative assessments across sample groups. Together, these analyses lead to a holistic exploration of the soil phytohormonome. Our method enables further research into soil metabolite and phytohormone dynamics, helping to elucidate future paradigms of community resilience in the understudied ecosystem of the soil environment.

## Abbreviations

2IP: N6-isopentenyladenine
ABA: abscisic acid
BA: N6-benzyladenine
GA: gibberellic acid
IAA: indole-3-acetic acid
IBA: indole-3-butyric acid
JA: jasmonic acid
KAR1: Karrikin 1
KAR2: Karrikin 2
KAR3: Karrikin 3
KAR4: Karrikin 4
Mel: melatonin
PGR: plant growth regulator
SA: salicylic acid
SPE: solid phase extraction
Trp: tryptophan
UPLC-qTOF-MS/MS: ultra-performance liquid chromatography quadrupole time-of-flight tandem mass-spectrometry
Zea: zeatin

## Supplementary Information

The following supplementary information are available:

Table S1: Mean quantity of analytes detected during Youden analysis.

Table S2: Analyte target list for targeted quantitation of phytohormones.

Table S3: Target list used for untargeted analysis of soils.

Table S4: Significant annotations from untargeted analysis and post-hoc testing after one-way ANOVA.

## Author Contributions

SL: conceptualization, methods, investigation, formal analysis, data curation, writing – draft, writing – review and editing, visualization. RZ: conceptualization, investigation, writing – review and editing. CF: supervision, resources, funding acquisition. JC: conceptualization, writing – review and editing, resources, funding acquisition LE: conceptualization, writing – review and editing, resources, supervision, project administration, funding acquisition.

## Data Availability

Metabolomics data used in this article are available at The Metabolomics Workbench (www.metabolomicsworkbench.org) doi:xxx. A full step-by-step protocol for the extraction methods described here has been deposited at protocols.io and it is freely available at https://dx.doi.org/10.17504/protocols.io.eq2lyoy6wgx9/v1

