## Supplemental Table for "Validated phytohormone analysis with paired untargeted workflow enables new insights into soil phytohormonome"

*Table S 1. Mean quantity of analytes (pg/g moisture-corrected dry weight) following Youden analysis with an 8-run, 6 factor fractional factorial design.*

| **Factor** | **Levels** | | **ABA**[1] | **IAA**[1] | **IBA**[1] | **JA**[1] | **KAR1**[1] | **KAR2**[1] | **KAR3**[1] | **TRP**[1] | **ZEA**[1] | **GA3**[1] | **SA**[1] |
|---|---|---|---|---|---|---|---|---|---|---|---|---|---|
| ACID | Formic acid | Level (+1) | 28.70 | 33.15 | 45.24 | 34.82 | 0.54 | 1.60 | 0.00 | 22.07 | 4.86 | 130.21 | 80.92 |
| | Acetic acid | Level (-1) | 21.44 | 35.25 | 47.34 | 29.87 | 2.03 | 0.00 | 0.91 | 25.83 | 4.34 | 153.54 | 93.77 |
| SALT | 1 M NaCl | Level (+1) | 30.87 | 33.26 | 46.08 | 34.37 | 0.00 | 0.00 | 0.00 | 20.14 | 1.45 | 141.00 | 111.09 |
| | None | Level (-1) | 22.70 | 35.03 | 46.31 | 32.01 | 3.09 | 2.62 | 0.80 | 23.94 | 6.34 | 150.11 | 75.67 |
| BASE | 0.1N NaOH | Level (+1) | 40.66 | 36.52 | 45.93 | 41.85 | 3.09 | 0.47 | 0.45 | 19.27 | 0.72 | 161.29 | 137.97 |
| | 0.01N NaOH | Level (-1) | 12.91 | 31.78 | 46.46 | 24.53 | 0.00 | 1.13 | 0.29 | 24.82 | 7.07 | 129.82 | 48.80 |
| ORDER | Acid, Base | Level (+1) | 37.11 | 35.28 | 46.15 | 42.11 | 2.54 | 0.00 | 0.40 | 13.32 | 0.00 | 164.00 | 110.81 |
| | Base, Acid | Level (-1) | 16.46 | 33.02 | 46.24 | 24.27 | 0.54 | 2.62 | 0.34 | 30.76 | 8.40 | 127.11 | 75.95 |
| TIME | Increased Vortex | Level (+1) | 32.57 | 36.30 | 45.58 | 41.25 | 0.54 | 0.47 | 0.00 | 28.77 | 3.65 | 154.51 | 106.74 |
| | Increased Sonicate | Level (-1) | 20.99 | 32.00 | 46.81 | 25.13 | 2.54 | 1.13 | 0.85 | 15.32 | 4.14 | 136.60 | 80.02 |
| WET | Prewet, 5 mL 1M NaCl | Level (+1) | 30.89 | 32.23 | 45.65 | 32.52 | 0.00 | 0.00 | 0.23 | 17.28 | 3.53 | 146.20 | 114.17 |
| | None | Level (-1) | 22.67 | 36.07 | 46.74 | 33.85 | 3.09 | 1.93 | 0.51 | 26.81 | 4.26 | 144.91 | 72.59 |

[1]ABA: Abscisic acid; IAA: indole-3-acetic acid; IBA: indole-3-butyric acid; JA: jasmonic acid; KAR1 – 3: karrikins 1-3; Trp: tryptophan; Zea: zeatin; GA3: gibberellic acid A3; SA: salicylic acid.

*Table S 2: Analyte target list for targeted quantification of analytes. Underlined ions indicate they are used in quantification, bolded ions indicate [M+H]$^+$ or [M-H]$^-$ pseudomolecular ions.*

| Analyte[1] | Ionization | Target Ions | RT[2] | Ion Ratio | Area Threshold | Height Threshold |
|---|---|---|---|---|---|---|
| **ABA** | NEG (-) | 527.2498, 264.1323, **263.1289**, 219.1338, 153.0822 | 10.1 | 0.166 | 3000 | 1500 |
| **IAA** | POS (+) | 177.0738, **176.0706**, 130.0665 | 9.61 | 0.112 | 500 | 500 |
| **IBA** | POS (+) | 205.1051, **204.1019**, 186.0918, 130.0665 | 10.72 | 0.133 | 500 | 500 |
| **JA** | NEG (-) | 210.1217, **209.1183**, 165.1253, 103.9185 | 10.8 | 0.133 | 3000 | 1500 |
| **KAR1** | POS (+) | 152.0424, **151.039** | 9 | 0.088 | 1500 | 500 |
| **KAR2** | POS (+) | 138.0267, **137.0233** | 5.96 | 0.072 | 1500 | 500 |
| **KAR3** | POS (+) | 166.058, **165.0546** | 10.23 | 0.097 | 1500 | 500 |
| **KAR4** | POS (+) | 166.058, **165.0546** | 10 | 0.096 | 1500 | 500 |
| **Mel** | POS (+) | 234.1315, **233.1285**, 174.0916 | 9.52 | 0.136 | 500 | 500 |
| **BA** | POS (+) | 227.1113, **226.1087**, 91.0546 | 8.72 | 0.13 | 500 | 500 |
| **2iP** | POS (+) | 205.1269, **204.1244**, 136.0625 | 8.52 | 0.108 | 500 | 500 |
| **5HT** | POS (+) | 178.1052, **177.1022**, 160.0761 | 2.84 | 0.14 | 1500 | 500 |
| **Trp** | POS (+) | 206.1002, **205.0972**, 188.0709 | 5.52 | 0.118 | 1500 | 500 |
| **Zea** | POS (+) | 221.1218, **220.1193**, 136.0633 | 4.96 | 0.114 | 1500 | 500 |
| **GA3** | NEG (-) | 363.1338, 346.1378, **345.1344**, 283.1267, 143.0838 | 8.6 | 0.21 | 3000 | 1500 |
| **SA** | NEG (-) | 236.9438, 174.0511, 138.0278, **137.0244**, 93.0331 | 9.5 | 0.077 | 3000 | 1500 |

[1]ABA: Abscisic acid; IAA: indole-3-acetic acid; IBA: indole-3-butyric acid; JA: jasmonic acid; KAR1 – 4: karrikins 1-4; BA: N6-benzyladenine; 2IP: N6-isopentenyladenine; 5HT: serotonin; Trp: tryptophan; Zea: zeatin; GA3: gibberellic acid A3; SA: salicylic acid.

[2]RT: Retention time, reported in minutes.

*Table S 3: Target list for untargeted metabolomics.*

| Name | Formula | Main Positive Adduct | [M+H]+ | Main Negative Adduct | [M-H]- |
|---|---|---|---|---|---|
| 12-hydroxy-jasmonic acid | C18H28O9 | [M+H]+ | 389.18063 | [M-H]- | 387.166 |
| 2,3-dihydroxyindole | C8H7NO2 | [M+H]+ | 150.055 | [M-H]- | 148.0404 |
| 24-epicastasterone | C28H48O5 | [M+H]+ | 465.35748 | [M-H]- | 463.3429 |
| 28-norcastasterone | C27H46O5 | [M+H]+ | 451.34183 | [M-H]- | 449.3272 |
| 2-amino-3-carboxymuconic acid-semialdehyde | C7H7NO5 | [M+H]+ | 186.03973 | [M-H]- | 184.0251 |
| 2-aminomuconate | C6H7NO4 | [M+H]+ | 158.0448 | [M-H]- | 156.0302 |
| 2-Aminomuconic semialdehyde | C6H7NO3 | [M+H]+ | 142.04989 | [M-H]- | 140.0353 |
| 2-aminophenol | C6H7NO | [M+H]+ | 110.0601 | [M-H]- | 108.0455 |
| 2-formamino-benzoylacetate | C10H9NO4 | [M+H]+ | 208.0605 | [M-H]- | 206.0459 |
| 2-formylamino-benzaldehyde | C7H7NO | [M+H]+ | 150.055 | [M-H]- | 148.0404 |
| 2-hydroxymelatonin | C13H18N2O3 | [M+H]+ | 251.3053 | [M-H]- | 249.2907 |
| 2-methylthio-cis-zeatin | C11H15N5OS | [M+H]+ | 266.10703 | [M-H]- | 264.0924 |
| 2-methylthio-cis-zeatin riboside | C16H23N5O5S | [M+H]+ | 398.14929 | [M-H]- | 396.1347 |
| 2-methylthio-isopentenyladenine | C11H15N5S | [M+H]+ | 250.11212 | [M-H]- | 248.0975 |
| 2-methylthio-isopentenyladenosine | C16H23N5O4S | [M+H]+ | 382.15438 | [M-H]- | 380.1398 |
| 2-Oxoadipic acid | C6H8O5 | [M+H]+ | 161.0445 | [M-H]- | 159.0299 |
| 2-oxoindole-3-acetate | C10H9NO3 | [M+H]+ | 192.0655 | [M-H]- | 190.0509 |
| 2-oxo-indole-3-acetic acid | C10H9NO3 | [M+H]+ | 192.19066 | [M-H]- | 190.1761 |
| 3-Hydroxyanthranilic acid | C7H7NO3 | [M+H]+ | 153.04262 | [M-H]- | 151.028 |
| 3-hydroxy-butanoyl-CoA | C25H42N7O18P3S | [M+H]+ | 854.1593 | [M-H]- | 852.1447 |
| 3-hydroxykynurenine | C10H12N2O4 | [M+H]+ | 225.08701 | [M-H]- | 223.0724 |
| 3-indole glycol-aldehyde | C10H9NO2 | [M+H]+ | 176.0706 | [M-H]- | 174.056 |
| 3-methoxy-anthranilate | C8H9NO3 | [M+H]+ | 168.0655 | [M-H]- | 166.0509 |
| 3-methyl-dioxy indole | C9H9NO2 | [M+H]+ | 164.0706 | [M-H]- | 162.056 |
| 3-methyl-indolepyruvate | C12H11NO3 | [M+H]+ | 218.0863 | [M-H]- | 216.0717 |
| 3-oxo-2-(2-(Z)-pentenyl)cyclopentane-1-butyric acid | C14H22O3 | [M+H]+ | 239.1642 | [M-H]- | 237.1496 |
| 3-oxo-2-(2-(Z)-pentenyl)cyclopentane-1-hexanoic acid | C16H26O3 | [M+H]+ | 267.1955 | [M-H]- | 265.1809 |
| 4-(2-amino-5-hydroxyphenyl)-2,4-dioxobutanoate | C10H9NO5 | [M+H]+ | 224.0554 | [M-H]- | 222.0408 |

| Name | Formula | Main Positive Adduct | [M+H]+ | Main Negative Adduct | [M-H]- |
|---|---|---|---|---|---|
| 4,6-dihydroxyquinoline | C9H7NO2 | [M+H]+ | 162.055 | [M-H]- | 160.0404 |
| 4,8-dihydroxyquinoline | C9H7NO2 | [M+H]+ | 162.055 | [M-H]- | 160.0404 |
| 4-hydroxymelatonin | C13H16N2O3 | [M+H]+ | 249.12339 | [M-H]- | 247.1088 |
| 5-(2'-carboxyethyl)-4,6-dihydroxypicolinate | C9H9NO6 | [M+H]+ | 228.0503 | [M-H]- | 226.0357 |
| 5-(2'-formylethyl)-4,6-dihydroxypicolinate | C9H9NO5 | [M+H]+ | 212.0554 | [M-H]- | 210.0408 |
| 5-(3'-carboxy-3'-oxopropenyl)-4,6-dihydroxypicolinate | C10H7NO7 | [M+H]+ | 254.0296 | [M-H]- | 252.015 |
| 5-(3'-carboxy-3'-oxopropyl)-4,6-dihydroxypicolinate | C10H9NO7 | [M+H]+ | 256.0452 | [M-H]- | 254.0306 |
| 5-deoxystrigol | C19H22O5 | [M+H]+ | 331.154 | [M-H]- | 329.1394 |
| 5-hydroxy tryptophan | C11H12N2O3 | [M+H]+ | 221.09209 | [M-H]- | 219.0775 |
| 5-hydroxyindole-acetaldehyde | C10H9NO2 | [M+H]+ | 176.0706 | [M-H]- | 174.056 |
| 5-hydroxy-indoleacetate | C10H9NO3 | [M+H]+ | 192.0655 | [M-H]- | 190.0509 |
| 5-hydroxyindole-acetylglycine | C12H12N2O4 | [M+H]+ | 249.087 | [M-H]- | 247.0724 |
| 5-hydroxy-indolepyruvate | C11H9NO4 | [M+H]+ | 220.0605 | [M-H]- | 218.0459 |
| 5-hydroxy-kynureamine | C10H12N2O4 | [M+H]+ | 181.0972 | [M-H]- | 179.0826 |
| 5-Hydroxykynurenine | C10H12N2O4 | [M+H]+ | 181.0972 | [M-H]- | 179.0826 |
| 5-hydroxy-N-formylkynurenine | C11H12N2O5 | [M+H]+ | 253.0819 | [M-H]- | 251.0673 |
| 5-Methoxy-1H-indole-3-carbaldehyde | C10H9NO2 | [M+H]+ | 176.07063 | [M-H]- | 174.056 |
| 5-methoxyindole-3 acetic acid | C11H11NO3 | [M+H]+ | 206.08119 | [M-H]- | 204.0666 |
| 5-methoxy-indoleacetate | C11H11NO3 | [M+H]+ | 206.0812 | [M-H]- | 204.0666 |
| 5-methoxytryptamine | C11H14N2O | [M+H]+ | 191.11791 | [M-H]- | 189.1033 |
| 5-methoxytryptamine | C11H14N2O | [M+H]+ | 191.1179 | [M-H]- | 189.1033 |
| 5-methoxytryptophol | C11H13NO2 | [M+H]+ | 192.10193 | [M-H]- | 190.0873 |
| 6-hydroxy-kynurenate | C10H7NO4 | [M+H]+ | 206.0448 | [M-H]- | 204.0302 |
| 6-hydroxymelatonin | C13H16N2O3 | [M+H]+ | 249.12339 | [M-H]- | 247.1088 |
| 7,8-dihydroxykynurenate | C10H7NO5 | [M+H]+ | 222.0397 | [M-H]- | 220.0251 |
| 7,8-hidydro-7,8-dihydroxykynurenate | C10H9NO5 | [M+H]+ | 224.0554 | [M-H]- | 222.0408 |
| 7-hydroxy-ABA | C15H20O5 | [M+H]+ | 281.13837 | [M-H]- | 279.1238 |
| 7-hydroxymelatonin | C13H16N2O3 | [M+H]+ | 249.12339 | [M-H]- | 247.1088 |
| 8-methoxy-kynurenate | C11H9NO4 | [M+H]+ | 220.0605 | [M-H]- | 218.0459 |
| 9,10-dihydrojasmonic acid | C12H20O3 | [M+H]+ | 213.14854 | [M-H]- | 211.1339 |
| 9-hydroxy-ABA | C15H20O5 | [M+H]+ | 281.13837 | [M-H]- | 279.1238 |

| **Name** | **Formula** | **Main Positive Adduct** | **[M+H]+** | **Main Negative Adduct** | **[M-H]-** |
|---|---|---|---|---|---|
| Abscisic acid | C15H20O4 | [M+H]+ | 265.14346 | [M-H]- | 263.1289 |
| acetoacetyl coa | C25H40N7O18P3S | [M+H]+ | 852.1436 | [M-H]- | 850.129 |
| acetyl coa | C23H38N7O17P3S | [M+H]+ | 810.1331 | [M-H]- | 808.1185 |
| acetylindoxyl | C10H9NO2 | [M+H]+ | 176.0706 | [M-H]- | 174.056 |
| Aminomethylphosphonic Acid | CH6NO3P | [M+H]+ | 112.01583 | [M-H]- | 110.0012 |
| aminomuconate | C6H7NO4 | [M+H]+ | 157.03698 | [M-H]- | 155.0224 |
| aniline | C6H7N | [M+H]+ | | [M-H]- | |
| anthranilate | C7H7NO2 | [M+H]+ | 137.04771 | [M-H]- | 135.0331 |
| anthranilic acid | C7H7NO2 | [M+H]+ | 138.05498 | [M-H]- | 136.0404 |
| betalamic acid | C9H9NO5 | [M+H]+ | 212.05537 | [M-H]- | 210.0408 |
| brassinolide | C28H48O6 | [M+H]+ | 481.35239 | [M-H]- | 479.3378 |
| bufotenine | C12H16N2O | [M+H]+ | 205.13356 | [M-H]- | 203.119 |
| Cadaverine | C5H14N2 | [M+H]+ | 103.123 | [M-H]- | 101.1084 |
| caldohexamine | C15H38N6 | [M+H]+ | 303.3231 | [M-H]- | 301.3085 |
| caldopentamine | C12H31N5 | [M+H]+ | 246.26525 | [M-H]- | 244.2507 |
| castasterone | C28H48O5 | [M+H]+ | 465.35748 | [M-H]- | 463.3429 |
| cinnamoyl serotonin | C19H18N2O2 | [M+H]+ | 307.14413 | [M-H]- | 305.1295 |
| cinnavalininate | C14H8N2O6 | [M+H]+ | 301.0455 | [M-H]- | 299.0309 |
| cis-12-oxo-phytodienoic acid | C18H28O3 | [M+H]+ | 293.21115 | [M-H]- | 291.1966 |
| crotonyl-coA | C25H40N7O17P3S | [M+H]+ | 836.1487 | [M-H]- | 834.1341 |
| cyclic (3)-hydroxymelatonin | C13H16N2O3 | [M+H]+ | 247.10774 | [M-H]- | 245.0931 |
| cyclic-melatonin | C13H14N2O2 | [M+H]+ | 231.11283 | [M-H]- | 229.0982 |
| cyclo-dopa | C9H9NO4 | [M+H]+ | 196.06046 | [M-H]- | 194.0459 |
| cyclo-dopa-5-O-glucoside | C15H19NO9 | [M+H]+ | 358.11328 | [M-H]- | 356.0987 |
| dihydrophaseic acid | C15H22O5 | [M+H]+ | 283.15402 | [M-H]- | 281.1394 |
| dihydrozeatin | C10H15N5O | [M+H]+ | 222.13496 | [M-H]- | 220.1204 |
| dihydrozeatin riboside-O-glucoside | C21H33N5O10 | [M+H]+ | 516.23005 | [M-H]- | 514.2155 |
| dihydrozeatin-glucoside | C16H25N5O6 | [M+H]+ | 384.18778 | [M-H]- | 382.1732 |
| dinor-12-oxo-phytodienoic acid | C18H24O3 | [M+H]+ | 265.18005 | [M-H]- | 263.1655 |
| dolicholide | C28H46O6 | [M+H]+ | 479.33674 | [M-H]- | 477.3221 |
| dolichosterone | C28H46O5 | [M+H]+ | 463.34183 | [M-H]- | 461.3272 |

| Name | Formula | Main Positive Adduct | [M+H]+ | Main Negative Adduct | [M-H]- |
|---|---|---|---|---|---|
| dopamine | C8H11NO2 | [M+H]+ | 154.08628 | [M-H]- | 152.0717 |
| dopaquinone | C9H9NO4 | [M+H]+ | 196.06046 | [M-H]- | 194.0459 |
| Epinephrine | C9H13NO3 | [M+H]+ | 184.09684 | [M-H]- | 182.0822 |
| feruloyl serotonin | C20H20N2O4 | [M+H]+ | 353.14961 | [M-H]- | 351.135 |
| formyl-5-hydroxy-kynureamine | C10H12N2O3 | [M+H]+ | 209.0921 | [M-H]- | 207.0775 |
| formylanthranilate | C8H7NO3 | [M+H]+ | 165.04207 | [M-H]- | 163.0275 |
| formylkynurenine | C11H12N2O4 | [M+H]+ | 237.08701 | [M-H]- | 235.0724 |
| formyl-n-acetyl-5-methoxykynureamine | C13H16N2O4 | [M+H]+ | 265.1183 | [M-H]- | 263.1037 |
| gamma-oxalo-crotonate | C6H6O5 | [M+H]+ | 159.0288 | [M-H]- | 157.0142 |
| Gibberellin A1 | C19H24O6 | [M+H]+ | 349.16459 | [M-H]- | 347.15 |
| Gibberellin A15 | C21H28O4 | [M+H]+ | 345.20606 | [M-H]- | 343.1915 |
| Gibberellin A19 | C20H26O6 | [M+H]+ | 363.18024 | [M-H]- | 361.1656 |
| Gibberellin A24 | C20H26O5 | [M+H]+ | 347.18532 | [M-H]- | 345.1707 |
| Gibberellin A29 | C19H24O6 | [M+H]+ | 349.16459 | [M-H]- | 347.15 |
| Gibberellin A3 | C19H22O6 | [M+H]+ | 347.14894 | [M-H]- | 345.1343 |
| Gibberellin A34 | C19H24O6 | [M+H]+ | 349.16459 | [M-H]- | 347.15 |
| Gibberellin A4 | C19H24O5 | [M+H]+ | 333.16967 | [M-H]- | 331.1551 |
| Gibberellin A44 | C20H26O5 | [M+H]+ | 347.18532 | [M-H]- | 345.1707 |
| Gibberellin A5 | C19H22O5 | [M+H]+ | 331.15402 | [M-H]- | 329.1394 |
| Gibberellin A51 | C19H24O5 | [M+H]+ | 333.16967 | [M-H]- | 331.1551 |
| Gibberellin A53 | C20H28O5 | [M+H]+ | 349.20097 | [M-H]- | 347.1864 |
| Gibberellin A6 | C19H22O6 | [M+H]+ | 347.14894 | [M-H]- | 345.1343 |
| Gibberellin A8 | C19H24O7 | [M+H]+ | 365.1595 | [M-H]- | 363.1449 |
| glucobrassicin | C16H20N2O9S2 | [M+H]+ | 449.0683 | [M-H]- | 447.0537 |
| glutaryl CoA | C26H42N7O19P3S | [M+H]+ | 882.1542 | [M-H]- | 880.1396 |
| Glyphosate | C3H8NO5P | [M+H]+ | 170.02131 | [M-H]- | 168.0067 |
| homobrassinolide | C29H50O6 | [M+H]+ | 495.36804 | [M-H]- | 493.3534 |
| homocastasterone | C29H50O5 | [M+H]+ | 479.37313 | [M-H]- | 477.3585 |
| homodolicholide | C29H48O6 | [M+H]+ | 493.35239 | [M-H]- | 491.3378 |
| homodolichosterone | C29H48O5 | [M+H]+ | 477.35748 | [M-H]- | 475.3429 |
| hydroxymelatonin | C13H16N2O3 | [M+H]+ | 249.1234 | [M-H]- | 247.1088 |

| Name | Formula | Main Positive Adduct | [M+H]+ | Main Negative Adduct | [M-H]- |
|---|---|---|---|---|---|
| IAA alanine | C13H14N2O3 | [M+H]+ | 247.10774 | [M-H]- | 245.0931 |
| IAA aspartate | C20H22N2O10 | [M+H]+ | 451.13475 | [M-H]- | 449.1202 |
| IAA glutamate | C21H24N2O10 | [M+H]+ | 465.1504 | [M-H]- | 463.1358 |
| IAA glycine | C12H12N2O3 | [M+H]+ | 233.24258 | [M-H]- | 231.228 |
| IAA leucine | C16H20N2O3 | [M+H]+ | 289.15469 | [M-H]- | 287.1401 |
| IAA phenylalanine | C19H18N2O3 | [M+H]+ | 323.13904 | [M-H]- | 321.1244 |
| IAA valine | C15H17N2O4 | [M+H]+ | 290.12613 | [M-H]- | 288.1115 |
| indol-3-yl-lactate | C11H11NO3 | [M+H]+ | 206.0812 | [M-H]- | 204.0666 |
| indole | C8H7N | [M+H]+ | 118.0651 | [M-H]- | 116.0505 |
| indole-3-acetaldehyde | C10H9NO | [M+H]+ | 160.0757 | [M-H]- | 158.0611 |
| indole-3-acetaldoxime | C10H10N2O | [M+H]+ | 175.0866 | [M-H]- | 173.072 |
| indole-3-acetamide | C10H10N2O | [M+H]+ | 525.27729 | [M-H]- | 523.2627 |
| indole-3-acetic acid | C10H9NO2 | [M+H]+ | 176.07063 | [M-H]- | 174.056 |
| indole-3-acetonitrile | C10H8N2 | [M+H]+ | 157.07605 | [M-H]- | 155.0615 |
| indole-3-butyric acid | C12H13NO2 | [M+H]+ | 204.10193 | [M-H]- | 202.0873 |
| indole-3-ethanol | C10H11NO | [M+H]+ | 162.0914 | [M-H]- | 160.0768 |
| indole-3-thiohydroximate | C10H10N2OS | [M+H]+ | 207.0587 | [M-H]- | 205.0441 |
| indolepyruvate | C11H9NO3 | [M+H]+ | 204.0655 | [M-H]- | 202.0509 |
| indolylmethyldesulfoglucosinolate | C16H20N2O6S | [M+H]+ | 369.1115 | [M-H]- | 367.0969 |
| indoxyl | C8H7NO | [M+H]+ | 134.0601 | [M-H]- | 132.0455 |
| isophenoxazine | C12H8N2O2 | [M+H]+ | 213.0659 | [M-H]- | 211.0513 |
| JA-isoleucine | C18H28NO4 | [M+H]+ | 324.21696 | [M-H]- | 322.2024 |
| JA-pheylalanine | C20H26NO5 | [M+H]+ | 358.20131 | [M-H]- | 356.1867 |
| jasmonic acid | C12H18O3 | [M+H]+ | 211.13289 | [M-H]- | 209.1183 |
| JA-tryptophan | C22H27N2O5 | [M+H]+ | 397.21221 | [M-H]- | 395.1976 |
| JA-valine | C16H26NO5 | [M+H]+ | 310.20131 | [M-H]- | 308.1867 |
| Karrikin 11 | C8H6O3 | [M+H]+ | 173.0445 | [M-H]- | 171.0299 |
| Karrikin 11 | C7H8O5 | [M+H]+ | 151.03899 | [M-H]- | 149.0244 |
| Karrikin 2 | C7H4O3 | [M+H]+ | 137.0233 | [M-H]- | 135.0087 |
| Karrikin 3 | C9H8O3 | [M+H]+ | 165.0546 | [M-H]- | 163.04 |
| Karrikin 4 | C9H8O4 | [M+H]+ | 165.0546 | [M-H]- | 163.04 |

| Name | Formula | Main Positive Adduct | [M+H]+ | Main Negative Adduct | [M-H]- |
|---|---|---|---|---|---|
| Karrikin 5 | C8H6O3 | [M+H]+ | 151.039 | [M-H]- | 149.0244 |
| Karrikin 6 | C8H6O4 | [M+H]+ | 151.039 | [M-H]- | 149.0244 |
| kinetin | C10H9N5O | [M+H]+ | 216.08801 | [M-H]- | 214.0734 |
| kinetin-9-glucoside | C16H19N5O6 | [M+H]+ | 378.14084 | [M-H]- | 376.1262 |
| kinetin-riboside | C15H17N5O5 | [M+H]+ | 348.13027 | [M-H]- | 346.1157 |
| kynuramine | C9H12N2O | [M+H]+ | 165.10226 | [M-H]- | 163.0877 |
| kynurenate | C10H7NO3 | [M+H]+ | 190.0499 | [M-H]- | 188.0353 |
| kynurenine | C10H12N2O3 | [M+H]+ | 209.09209 | [M-H]- | 207.0775 |
| levodopa | C9H11NO4 | [M+H]+ | 198.07611 | [M-H]- | 196.0615 |
| MEL-Ala | C16H21N3O3 | [M+H]+ | 304.16559 | [M-H]- | 302.151 |
| MEL-Arg | C19H36N6O3 | [M+H]+ | 389.22959 | [M-H]- | 387.215 |
| MEL-Asn | C17H22N4O4 | [M+H]+ | 347.17141 | [M-H]- | 345.1568 |
| MEL-Asp | C17H21N3O5 | [M+H]+ | 348.15542 | [M-H]- | 346.1408 |
| melatonin | C13H16N2O2 | [M+H]+ | 233.12848 | [M-H]- | 231.1139 |
| MEL-Cys | C16H21N3O3S | [M+H]+ | 336.13766 | [M-H]- | 334.1231 |
| MEL-Gln | C18H24N4O4 | [M+H]+ | 361.18706 | [M-H]- | 359.1725 |
| MEL-Glu | C18H23N3O5 | [M+H]+ | 362.17107 | [M-H]- | 360.1565 |
| MEL-Gly | C15H19N3O3 | [M+H]+ | 290.14994 | [M-H]- | 288.1353 |
| MEL-His | C19H23N5O3 | [M+H]+ | 370.18739 | [M-H]- | 368.1728 |
| MEL-Ile | C19H27N3O3 | [M+H]+ | 346.21254 | [M-H]- | 344.1979 |
| MEL-Leu | C19H27N3O3 | [M+H]+ | 346.21254 | [M-H]- | 344.1979 |
| MEL-Lys | C19H28N4O3 | [M+H]+ | 361.22344 | [M-H]- | 359.2088 |
| MEL-Met | C18H25N3O3S | [M+H]+ | 364.16896 | [M-H]- | 362.1544 |
| MEL-Phe | C22H25N3O3 | [M+H]+ | 380.19689 | [M-H]- | 378.1823 |
| MEL-Pro | C18H23N3O3 | [M+H]+ | 330.18124 | [M-H]- | 328.1666 |
| MEL-Ser | C16H12N3O4 | [M+H]+ | 320.16051 | [M-H]- | 318.1459 |
| MEL-Thr | C17H23N3O4 | [M+H]+ | 334.17616 | [M-H]- | 332.1616 |
| MEL-Trp | C24H26N4O3 | [M+H]+ | 419.20779 | [M-H]- | 417.1932 |
| MEL-Tyr | C22H25N3O4 | [M+H]+ | 396.19181 | [M-H]- | 394.1772 |
| MEL-Val | C18H25N3O3 | [M+H]+ | 332.19689 | [M-H]- | 330.1823 |
| meta-topolin | C12H11N5O | [M+H]+ | 242.10366 | [M-H]- | 240.0891 |

| Name | Formula | Main Positive Adduct | [M+H]+ | Main Negative Adduct | [M-H]- |
|---|---|---|---|---|---|
| meta-topolin riboside | C17H19N5O7 | [M+H]+ | 374.14592 | [M-H]- | 372.1313 |
| meta-topolin-9-glucoside | C18H21N5O7 | [M+H]+ | 420.1514 | [M-H]- | 418.1368 |
| N6-benzyladenine | C12H11N5 | [M+H]+ | 226.10875 | [M-H]- | 224.0942 |
| N6-benzyladenine riboside | C12H11N5 | [M+H]+ | 358.151 | [M-H]- | 356.1364 |
| N6-benzyladenine-7,9-diglucoside | C24H33N5O10 | [M+H]+ | 552.23004 | [M-H]- | 550.2154 |
| N6-benzyladenine-7-glucoside | C18H21N5O5 | [M+H]+ | 388.16157 | [M-H]- | 386.147 |
| N6-benzyladenine-9-glucoside | C18H23N5O5 | [M+H]+ | 390.17722 | [M-H]- | 388.1626 |
| N6-isopentenyladenine | C23H29N5O9 | [M+H]+ | 204.1244 | [M-H]- | 202.1098 |
| N6-isopentenyladenine-7-glucoside | C10H13N5O | [M+H]+ | 366.17722 | [M-H]- | 364.1626 |
| N6-isopentenyladenosine | C16H23N5O5 | [M+H]+ | 336.16665 | [M-H]- | 334.1521 |
| N6-N2-formyl-5-methoxykynuramine | C15H22N5O7P | [M+H]+ | 265.11831 | [M-H]- | 263.1037 |
| N-acetylisatin | C10H7NO3 | [M+H]+ | 190.0499 | [M-H]- | 188.0353 |
| n-acetylserotonin | C12H14N2O2 | [M+H]+ | 219.1128 | [M-H]- | 217.0982 |
| N-caffeoyl Melatonin | C22H22N2O5 | [M+H]+ | 395.16017 | [M-H]- | 393.1456 |
| N-cinnamoyl Melatonin | C22H22N2O3 | [M+H]+ | 363.17034 | [M-H]- | 361.1557 |
| N-coumaroyl Melatonin | C22H22N2O4 | [M+H]+ | 379.16526 | [M-H]- | 377.1507 |
| neophaseic acid | C15H20O5 | [M+H]+ | 281.13837 | [M-H]- | 279.1238 |
| N-feruloyl Melatonin | C23H24N2O5 | [M+H]+ | 409.17582 | [M-H]- | 407.1612 |
| n-hydroxy-tryptamine | C15H19O5 | [M+H]+ | 177.1023 | [M-H]- | 175.0877 |
| nicotinic acid mononucleotide | C11H14NO9P | [M+H]+ | 337.05518 | [M-H]- | 335.0406 |
| n-methylserotonin | C11H14N2O | [M+H]+ | 191.1179 | [M-H]- | 189.1033 |
| N-methyltryptamine | C11H14N2 | [M+H]+ | 175.123 | [M-H]- | 173.1084 |
| norbrassinolide | C27H46O6 | [M+H]+ | 467.33674 | [M-H]- | 465.3221 |
| norepinephrine | C8H11NO3 | [M+H]+ | 170.08119 | [M-H]- | 168.0666 |
| N-sinapoyl Melatonin | C24H26N2O6 | [M+H]+ | 439.18639 | [M-H]- | 437.1718 |
| octopamine | C8H11NO2 | [M+H]+ | 154.08627 | [M-H]- | 152.0717 |
| olicholide | C28H46O6 | [M+H]+ | 479.33726 | [M-H]- | 477.3227 |
| orobanchol | C19H22O6 | [M+H]+ | 347.1489 | [M-H]- | 345.1343 |
| orobanchyl acetate | C21H24O7 | [M+H]+ | 389.1595 | [M-H]- | 387.1449 |
| Ortho-topolin | C12H11N5O | [M+H]+ | 242.10366 | [M-H]- | 240.0891 |
| Ortho-topolin riboside | C17H19N5O6 | [M+H]+ | 374.14592 | [M-H]- | 372.1313 |

| Name | Formula | Main Positive Adduct | [M+H]+ | Main Negative Adduct | [M-H]- |
|---|---|---|---|---|---|
| ortho-tropolin-9-glucoside | C18H21N5O8 | [M+H]+ | 420.1514 | [M-H]- | 418.1368 |
| para-topolin riboside | C17H19N5O5 | [M+H]+ | 374.14592 | [M-H]- | 372.1313 |
| Para-tropolin | C12H11N5O | [M+H]+ | 242.10366 | [M-H]- | 240.0891 |
| para-tropolin-9-glucoside | C18H21N5O9 | [M+H]+ | 420.1514 | [M-H]- | 418.1368 |
| phaseic acid | C15H20O5 | [M+H]+ | 281.13837 | [M-H]- | 279.1238 |
| picolinate | C6H5NO2 | [M+H]+ | 124.03933 | [M-H]- | 122.0247 |
| Putrescine | C4H12N2 | [M+H]+ | | [M-H]- | |
| quinolinate | C7H5NO4 | [M+H]+ | 173.04771 | [M-H]- | 171.0331 |
| quinolinic acid | C7H5NO4 | [M+H]+ | 168.02916 | [M-H]- | 166.0146 |
| Salicyclic acid | C7H6O3 | [M+H]+ | 139.03899 | [M-H]- | 137.0244 |
| serotonin | C10H12N2O | [M+H]+ | 177.10226 | [M-H]- | 175.0877 |
| skatole | C9H9N | [M+H]+ | 132.0808 | [M-H]- | 130.0662 |
| sorgolactone | C18H20O5 | [M+H]+ | 317.1384 | [M-H]- | 315.1238 |
| Spermidine | C7H19N3 | [M+H]+ | 146.1652 | [M-H]- | 144.1506 |
| Spermine | C10H26N4 | [M+H]+ | 203.22305 | [M-H]- | 201.2085 |
| strigol | C19H22O6 | [M+H]+ | 347.1489 | [M-H]- | 345.1343 |
| strigyl acetate | C21H24O7 | [M+H]+ | 389.1595 | [M-H]- | 387.1449 |
| teasterone | C28H48O4 | [M+H]+ | 449.36256 | [M-H]- | 447.348 |
| tetrakis(3-aminopropyl) ammonium | C12H32N5 | [M+H]+ | 247.27307 | [M-H]- | 245.2585 |
| Thidiazuron | C9H8N4OS | [M+H]+ | 221.04918 | [M-H]- | 219.0346 |
| Thidiazuron (dimer) | C18H16N8O2S2 | [M+H]+ | 441.09106 | [M-H]- | 439.0765 |
| Thidiazuron (tetramer) | C36H32N16O4S4 | [M+H]+ | 881.17482 | [M-H]- | 879.1602 |
| Thidiazuron (trimer) | C27H24N12O3S3 | [M+H]+ | 661.13294 | [M-H]- | 659.1183 |
| tris(3-aminopropyl) amine | C9H24N4 | [M+H]+ | 189.2074 | [M-H]- | 187.1928 |
| tryptamine | C10H12N2 | [M+H]+ | 161.10735 | [M-H]- | 159.0928 |
| tryptophan | C11H12N2O2 | [M+H]+ | 205.09718 | [M-H]- | 203.0826 |
| typhasterol | C28H48O4 | [M+H]+ | 449.36256 | [M-H]- | 447.348 |
| tyramine | C8H11NO | [M+H]+ | 138.09136 | [M-H]- | 136.0768 |
| Tyrosine | C9H11NO3 | [M+H]+ | 182.08119 | [M-H]- | 180.0666 |
| xanthurenate | C10H7NO4 | [M+H]+ | 206.0448 | [M-H]- | 204.0302 |
| zeatin | C10H13N5O | [M+H]+ | 220.11931 | [M-H]- | 218.1047 |

| Name | Formula | Main Positive Adduct | [M+H]+ | Main Negative Adduct | [M-H]- |
|---|---|---|---|---|---|
| zeatin riboside | C15H21N5O5 | [M+H]+ | 352.16157 | [M-H]- | 350.147 |
| zeatin riboside-O-glucoside | C21H31N5O10 | [M+H]+ | 514.21439 | [M-H]- | 512.1998 |
| zeatin-7-glucoside | C16H25N5O6 | [M+H]+ | 384.18778 | [M-H]- | 382.1732 |
| zeatin-9-glucoside | C16H23N5O6 | [M+H]+ | 382.17213 | [M-H]- | 380.1575 |
| zeatin-O-glucoside | C16H23N5O6 | [M+H]+ | 382.17213 | [M-H]- | 380.1575 |

*Table S 4 : Significant analytes identified through one-way Analysis of Variance between extracts of different soil types, using a false discovery rate cutoff (FDR) of 0.05. Analytes in bold indicate where annotation was made using target list by exact mass, remaining analytes were annotated by publicly available spectral library.*

| **Sample** | ***F* Statistic** | **Raw *p*-value** | **FDR** | **Incidence by Soil Type** | | |
|---|---|---|---|---|---|---|
| | | | | **Ag. Field** | **Grasslands** | **Greenhouse** |
| RT0.87_Adenosine | 6.4 | 0.0327 | 0.0434 | Present | Present | Present |
| RT0.88_Deoxyadenosine | 23.7 | 0.0014 | 0.0030 | Present | Absent | Present |
| RT0.88_Epicatechin | 4.1 | 0.0747 | 0.0897 | Present | Absent | Present |
| **RT10.09_dihydrophaseic acid** | 143.9 | < 0.0001 | < 0.0001 | Present | Present | Present |
| RT10.1_(9Z,12E)-15,16-dihydroxyoctadeca-9,12-dienoic acid | 1.5 | 0.2926 | 0.3109 | Present | Present | Present |
| **RT10.2_Gibberellin A53** | 12.2 | 0.0076 | 0.0124 | Present | Present | Present |
| RT10.41_4-Methoxycinnamic acid | 1.9 | 0.2333 | 0.2505 | Present | Present | Present |
| **RT10.49_cis-12-oxo-phytodienoic acid** | 10.9 | 0.0101 | 0.0151 | Present | Absent | Present |
| RT10.5_(5R)-5-hydroxy-1-(4-hydroxy-3-methoxyphenyl)tetradecan-3-one | 1.9 | 0.2278 | 0.2472 | Present | Present | Present |
| RT10.51_CGA62826 (2-[2,6-dimethylphenyl)-methoxyacetylamino]propionic acid, metalaxyl metabolite) | 497.4 | < 0.0001 | < 0.0001 | Absent | Absent | Present |
| RT10.53_(9Z,12E)-15,16-dihydroxyoctadeca-9,12-dienoic acid | 0.8 | 0.4779 | 0.4924 | Present | Present | Present |
| RT10.62_(1R,2R,4aS,6aS,6bR,10S,12aR,14bS)-1,8,10-trihydroxy-1,2,6a,6b,9,9,12a-heptamethyl-2,3,4,5,6,6a,7,8,8a,10,11,12,13,14b-tetradecahydropicene-4a-carboxylic acid | 46.3 | 0.0002 | 0.0007 | Present | Present | Present |
| RT10.69_8-hydroxy-5,8a-dimethyl-3-methylidene-3a,4,4a,8,9,9a-hexahydrobenzo[f][1]benzofuran-2,7-dione | 41.0 | 0.0003 | 0.0008 | Present | Present | Present |
| RT10.94_NVE_247.1330_17.3 (Anisoles) | 13.0 | 0.0066 | 0.0110 | Present | Present | Present |
| **RT10.96_9,10-dihydrojasmonic acid** | 20.0 | 0.0022 | 0.0046 | Present | Present | Present |
| RT11.01_1-(3,4-dihydroxyphenyl)-7-(4-hydroxyphenyl)heptan-3-one | 107.7 | < 0.0001 | 0.0001 | Present | Present | Present |
| RT11.15_h_128_methyl_1_testosterone | 10.3 | 0.0114 | 0.0165 | Present | Present | Present |
| RT11.16_Atractylenolide III | 962.55 | < 0.0001 | < 0.0001 | Present | Present | Present |
| RT11.16_h_92_oxabolone | 3.5 | 0.1004 | 0.1138 | Present | Present | Present |

| Sample | *F* Statistic | Raw *p*-value | FDR | Incidence by Soil Type | | |
|---|---|---|---|---|---|---|
| | | | | Ag. Field | Grassland | Greenhouse |
| RT11.24_9-hydroxy-1,4a-dimethyl-7-propan-2-yl-2,3,4,9,10,10a-hexahydrophenanthrene-1-carboxylic acid | 5.4 | 0.0456 | 0.0589 | Present | Present | Present |
| RT11.4_(1S,4aS,5R)-5-[2-(furan-3-yl)ethyl]-1,4a-dimethyl-6-methylidene-3,4,5,7,8,8a-hexahydro-2H-naphthalene-1-carboxylic acid | 4.6 | 0.0615 | 0.0747 | Present | Present | Present |
| RT11.41_(1S,4S,5R,10S,13S,17S,19S,20R)-10-hydroxy-4,5,9,9,13,19,20-heptamethyl-24-oxahexacyclo[15.5.2.0?,??.0?,??.0?,??.0?,??]tetracos-15-en-23-one | 27.6 | 0.0009 | 0.0021 | Absent | Present | Absent |
| RT11.48_Azoxystrobin (free acid) | 47.9 | 0.0002 | 0.0006 | Present | Absent | Present |
| **RT11.51_9,10-dihydrojasmonic acid** | 55.2 | 0.0001 | 0.0005 | Present | Present | Present |
| RT11.57_9-hydroxy-1,4a-dimethyl-7-propan-2-yl-2,3,4,9,10,10a-hexahydrophenanthrene-1-carboxylic acid | 15.0 | 0.0046 | 0.0083 | Present | Present | Present |
| RT11.62_(10E,15Z)-9,12,13-trihydroxyoctadeca-10,15-dienoic acid | 6.5 | 0.0318 | 0.0426 | Absent | Present | Present |
| RT11.68_Isobavachalcone | 34.9 | 0.0005 | 0.0012 | Present | Present | Present |
| RT11.69_?-linolenic acid | 11.1 | 0.0097 | 0.0148 | Present | Present | Present |
| **RT11.73_kynuramine** | 790.76 | < 0.0001 | < 0.0001 | Present | Absent | Absent |
| **RT11.83_cis-12-oxo-phytodienoic acid** | 94.1 | < 0.0001 | 0.0001 | Present | Absent | Present |
| RT11.85_4-acetyloxy-8-(3-oxo-2-pent-2-enylcyclopenten-1-yl)octanoic acid | 276.4 | < 0.0001 | < 0.0001 | Present | Present | Present |
| RT11.91_Pesticide4_Chlorantraniliprole | 214.6 | < 0.0001 | < 0.0001 | Present | Present | Absent |
| **RT12.01_cis-12-oxo-phytodienoic acid** | 415.6 | < 0.0001 | < 0.0001 | Present | Present | Present |
| RT12.25_Norethindrone | 44.8 | 0.0002 | 0.0007 | Present | Present | Present |
| RT12.3_dihydroalbocycline | 2.5 | 0.1619 | 0.1795 | Present | Present | Present |
| **RT12.3_dinor-12-oxo-phytodienoic acid** | 19.1 | 0.0025 | 0.0050 | Present | Absent | Present |
| RT12.37_(5R)-5-hydroxy-1-(4-hydroxy-3-methoxyphenyl)tetradecan-3-one | 5.2 | 0.0498 | 0.0635 | Present | Present | Present |
| RT12.45_albocycline | 15.2 | 0.0045 | 0.0081 | Present | Present | Present |
| RT12.47_Glycyrrhetinic Acid | 11.7 | 0.0085 | 0.0135 | Present | Present | Present |

| **Sample** | ***F* Statistic** | **Raw *p*-value** | **FDR** | **Incidence by Soil Type** | | |
|---|---|---|---|---|---|---|
| | | | | **Ag. Field** | **Grassland** | **Greenhouse** |
| RT12.5_Myclobutanil | 21129 | < 0.0001 | < 0.0001 | Absent | Absent | Present |
| RT12.5_Prothioconazole-desethio | 13.4 | 0.0061 | 0.0106 | Present | Present | Present |
| RT12.58_methyl (2E,4E,8E)-7,13-dihydroxy-4,8,12-trimethyltetradeca-2,4,8-trienoate | 14.9 | 0.0047 | 0.0083 | Present | Present | Absent |
| RT12.72_Nandrolone | 159.5 | < 0.0001 | < 0.0001 | Present | Present | Present |
| RT12.73_2,4-dihydroxyheptadec-16-ynyl acetate | 13.1 | 0.0065 | 0.0110 | Present | Present | Present |
| RT12.75_?-linolenic acid | 27.1 | 0.0010 | 0.0022 | Present | Present | Present |
| RT12.79_Methoxyfenozide | 395.8 | < 0.0001 | < 0.0001 | Present | Absent | Absent |
| RT12.8_Roccellic acid | 93.0 | < 0.0001 | 0.0001 | Present | Present | Absent |
| RT12.85_(9Z,12E)-15,16-dihydroxyoctadeca-9,12-dienoic acid | 6.9 | 0.0276 | 0.0381 | Present | Present | Present |
| **RT12.9_neophaseic acid** | 12.2 | 0.0077 | 0.0124 | Present | Present | Present |
| RT12_eudesmin | 77.2 | < 0.0001 | 0.0002 | Present | Present | Present |
| **RT13.01_cis-12-oxo-phytodienoic acid** | 0.4 | 0.6829 | 0.6897 | Present | Present | Present |
| RT13.03_NX2 toxin | 2284.7 | < 0.0001 | < 0.0001 | Absent | Absent | Present |
| RT13.1_(2S,3S,4S,8R,9S,13R,14R,15S,16R)-3,4,8,14,15-pentahydroxy-2,13,16-trimethyl-6-methylidene-10-oxatetracyclo[7.6.1.0?,?.0??,??]hexadecan-11-one | 520.0 | < 0.0001 | < 0.0001 | Absent | Absent | Present |
| RT13.13_Trifloxystrobin CGA 321113 | 761.2 | < 0.0001 | < 0.0001 | Absent | Absent | Present |
| RT13.18_(5R)-5-hydroxy-1-(4-hydroxy-3-methoxyphenyl)tetradecan-3-one | 2.1 | 0.2071 | 0.2271 | Present | Present | Present |
| RT13.22_h_61_17_epioxandrolone | 3.6 | 0.0950 | 0.1089 | Present | Present | Present |
| RT13.26_1,2-dihydroxyheptadec-16-en-4-yl acetate | 122.7 | < 0.0001 | 0.0001 | Present | Present | Present |
| RT13.26_5-[5-(acetyloxymethyl)-1,2,4a-trimethyl-7-oxo-3,4,8,8a-tetrahydro-2H-naphthalen-1-yl]-3-methylpentanoic acid | 1237.1 | < 0.0001 | < 0.0001 | Absent | Absent | Present |
| RT13.3_Norethindrone | 23.2 | 0.0015 | 0.0031 | Present | Present | Present |
| RT13.32_1-(3,4-dihydroxyphenyl)-7-(4-hydroxyphenyl)heptan-3-one | 212.2 | < 0.0001 | < 0.0001 | Present | Present | Present |

| Sample | *F* Statistic | Raw *p*-value | FDR | Incidence by Soil Type | | |
|---|---|---|---|---|---|---|
| | | | | Ag. Field | Grassland | Greenhouse |
| RT13.33_(1R,2R,5S,8R,14R,15R,16S)-16-hydroxy-1,2,14,17,17-pentamethyl-8-(prop-1-en-2-yl)pentacyclo[11.7.0.0?,??.0?,?.0??,??]icosane-5,15-dicarboxylic acid | 32.0 | 0.0006 | 0.0015 | Present | Present | Present |
| RT13.43_1,2-dihydroxyheptadec-16-en-4-yl acetate | 25.7 | 0.0011 | 0.0025 | Present | Present | Present |
| RT13.44_1,4a-dimethyl-9-oxo-7-propan-2-yl-3,4,10,10a-tetrahydro-2H-phenanthrene-1-carboxylic acid | 10.3 | 0.0115 | 0.0165 | Present | Present | Present |
| RT13.63_?-linolenic acid | 8.7 | 0.0169 | 0.0239 | Present | Present | Present |
| RT13.64_(E)-1-(2,4-dihydroxyphenyl)-3-[4-(3-methylbut-2-enoxy)phenyl]prop-2-en-1-one | 167.2 | < 0.0001 | < 0.0001 | Absent | Present | Absent |
| **RT13.68_3-methyl-dioxy indole** | 279.5 | < 0.0001 | < 0.0001 | Absent | Absent | Present |
| RT13.71_dehydroabietic acid | 17.2 | 0.0033 | 0.0062 | Present | Present | Present |
| RT13.79_h_2_etiocholanolone | 31.8 | 0.0006 | 0.0015 | Present | Absent | Present |
| RT13.84_Enoxolone | 0.4 | 0.6986 | 0.6986 | Present | Present | Present |
| RT13.9_?-linolenic acid | 3.8 | 0.0853 | 0.1000 | Present | Present | Present |
| RT14.07_(1S,4S,5R,10S,13S,17S,19S,20R)-10-hydroxy-4,5,9,9,13,19,20-heptamethyl-24-oxahexacyclo[15.5.2.0?,??.0?,??.0?,??.0?,??]tetracos-15-en-23-one | 5.1 | 0.0509 | 0.0641 | Present | Present | Present |
| RT14.08_Trifloxystrobin | 654.0 | < 0.0001 | < 0.0001 | Absent | Absent | Present |
| RT14.18_(E)-4-[(1R,2S,3S,4R,8aS)-2,3,4-trihydroxy-2,5,5,8a-tetramethyl-3,4,4a,6,7,8-hexahydro-1H-naphthalen-1-yl]but-3-en-2-one | 11.1 | 0.0096 | 0.0148 | Present | Present | Present |
| **RT14.2_cis-12-oxo-phytodienoic acid** | 10.6 | 0.0108 | 0.0159 | Present | Present | Present |
| RT14.33_h_82_tetrahydrogestrinone | 63.4 | < 0.0001 | 0.0003 | Present | Present | Present |
| RT14.58_Tetradecylsulfate | 0.7 | 0.5299 | 0.5405 | Present | Present | Present |
| **RT14.74_cis-12-oxo-phytodienoic acid** | 43.4 | 0.0003 | 0.0008 | Present | Present | Present |
| RT15.21_1,2-dihydroxyheptadec-16-en-4-yl acetate | 29.2 | 0.0008 | 0.0019 | Present | Present | Present |
| RT15.41_diosgenin | 5.0 | 0.0536 | 0.0666 | Present | Absent | Present |
| RT4.53_Homogentisic acid | 6.1 | 0.0362 | 0.0474 | Present | Present | Present |

| Sample | *F* Statistic | Raw *p*-value | FDR | Incidence by Soil Type | | |
|---|---|---|---|---|---|---|
| | | | | Ag. Field | Grassland | Greenhouse |
| RT6.77_Epicatechin | 16.6 | 0.0036 | 0.0067 | Present | Present | Present |
| **RT6.84_topolin riboside** | 99.5 | < 0.0001 | 0.0001 | Present | Present | Absent |
| **RT7.64_topolin** | 39.6 | 0.0003 | 0.0009 | Present | Present | Absent |
| RT7.95_fraxin | 17.4 | 0.0032 | 0.0061 | Present | Present | Present |
| RT8.07_Catechin | 1.3 | 0.3288 | 0.3458 | Present | Present | Present |
| RT8.24_(2R,3R,4S,5S,6R)-2-[3-[2-(4-hydroxy-3-methoxyphenyl)-3-(hydroxymethyl)-7-methoxy-2,3-dihydro-1-benzofuran-5-yl]propoxy]-6-(hydroxymethyl)oxane-3,4,5-triol | 51.9 | 0.0002 | 0.0005 | Present | Absent | Present |
| RT8.3_asperuloside | 77.3 | < 0.0001 | 0.0002 | Present | Absent | Absent |
| RT8.38_3-Hydroxy-4-methoxycinnamic acid (isoferulic acid) | 59.9 | 0.0001 | 0.0004 | Present | Present | Present |
| RT8.55_(2S,3S,4S,5R,6R)-6-(3-benzoyloxy-2-hydroxypropoxy)-3,4,5-trihydroxyoxane-2-carboxylic acid | 48.8 | 0.0002 | 0.0006 | Present | Present | Present |
| **RT8.55_topolin riboside** | 4.9 | 0.0557 | 0.0685 | Present | Present | Absent |
| RT8.59_P-Coumaric acid | 18.3 | 0.0028 | 0.0055 | Present | Present | Present |
| RT8.95_3-Hydroxy-4-methoxycinnamic acid (isoferulic acid) | 37.8 | 0.0004 | 0.0010 | Present | Present | Present |
| RT8.95_trans-Ferulic acid | 52.1 | 0.0002 | 0.0005 | Present | Present | Present |
| RT8_Fraxedin | 11.4 | 0.0091 | 0.0143 | Present | Present | Absent |
| RT9.27_3,4-Methylenedioxyamphetamine (MDA) | 1.1 | 0.3808 | 0.3964 | Present | Present | Present |
| RT9.3_N-Desmethyltramadol | 71.5 | < 0.0001 | 0.0003 | Present | Present | Present |
| RT9.37_DL-beta-Hydroxybutyric acid | 7.8 | 0.0213 | 0.0298 | Present | Present | Present |
| RT9.43_(2R,3S,4S,5R,6R)-2-(hydroxymethyl)-6-[[(2R,3S,4S,5R,6S)-3,4,5-trihydroxy-6-[4-[(E)-3-hydroxyprop-1-enyl]-2,6-dimethoxyphenoxy]oxan-2-yl]methoxy]oxane-3,4,5-triol | 3.9 | 0.0829 | 0.0983 | Present | Absent | Absent |
| **RT9.47_jasmonic acid** | 6.6 | 0.0302 | 0.0410 | Present | Present | Present |
| RT9.49_Unknown (carbon number 16); PlaSMA ID-1789 | 2.6 | 0.1538 | 0.1723 | Present | Present | Present |
| RT9.64_CGA62826 (2-[2,6-dimethylphenyl)-methoxyacetylamino]propionic acid, metalaxyl metabolite) | 41.3 | 0.0003 | 0.0008 | Absent | Absent | Present |
| RT9.73_4-[2-[(1R,4aS,5R,8aS)-6-hydroxy-5-(hydroxymethyl)-5,8a-dimethyl-2-methylidene-3,4,4a,6,7,8-hexahydro-1H-naphthalen-1-yl]ethyl]-2H-furan-5-one | 3.8 | 0.0871 | 0.1010 | Present | Present | Present |
